# Proteomic composition and mutual assembly of the C2a projection in vertebrate motile cilia

**DOI:** 10.64898/2026.01.24.701544

**Authors:** Qian Lyu, Qingchao Li, Jingrui Li, Jiajun Luo, Chunyu Liu, Shanshan Nai, Hongbin Liu, Xueliang Zhu, Ting Song, Min Liu, Huijie Zhao

## Abstract

The central apparatus of motile cilia, consisting of central microtubules and various protein projections, is essential for dictating the ciliary movement. Although three proteins (FAP65, FAP147, and FAP70) have been localized to the C2a projection in *Chlamydomonas reinhardtii*, the full protein composition and functional roles of the vertebrate C2a remain inadequately defined. Here, we use three knockout mouse models corresponding to their respective homologs (*Ccdc108*, *Mycbpap*, and *Cfap70*) to systematically investigate their functions in vertebrates. Notably, all three knockout strains exhibit distinct phenotypes related to primary ciliary dyskinesia (PCD), including hydrocephalus and sinusitis. The ciliary incorporation of CCDC108, MYCBPAP, and CFAP70 is essential for one another’s stability, with the loss of any single component triggering C2a collapse, which destabilizes the central pair microtubules and ultimately alters the ciliary movement pattern. Furthermore, we significantly expand the vertebrate C2a proteome by identifying ARMC3 and MYCBP as additional C2a components. Collectively, our findings illuminate the proteomic composition and strict physiological requirements of the vertebrate C2a projection, providing new insights into the molecular pathogenesis of PCD.

**Impact Statement:** Deficiency in any of the interdependent C2a proteins (CCDC108, MYCBPAP, and CFAP70) collapses this central pair microtubule-associated projection, disrupting vertebrate ciliary movement and causing primary ciliary dyskinesia phenotypes in mice.

## Introduction

Motile cilia and flagella are important motility-related organelles responsible for unicellular locomotion or generating fluid flow over the cell surfaces in vertebrates (*Lee and Ostrowski, 2021; Legendre et al., 2021; Lyu et al., 2024*). Typically, they exhibit a characteristic ‘9 + 2’ axoneme arrangement, featuring a central pair of microtubules (CP-MTs; C1 and C2) surrounded by nine peripheral doublet microtubules (DMTs). Multiple proteinous projections are formed on the CP-MTs, together forming the central apparatus (CA). The CA and another set of substructures on the DMTs, including dynein arms, radial spokes, and the nexin-dynein regulatory complex, cooperate to control proper ciliary motility (*Ishikawa, 2017; Lyu et al., 2024; Zhu, 2025*). Mutations in genes involved in these structures can affect ciliary motility and cause primary ciliary dyskinesia (PCD), a genetic disorder characterized by recurrent respiratory infections, situs inversus, infertility, and hydrocephalus in some cases (*Chen et al., 2026; Legendre et al., 2021; Reiter and Leroux, 2017; Song et al., 2025; Wallmeier et al., 2020*).

As an evolutionarily conserved structure, the CA primarily coordinates with radial spokes to regulate the default movement of DMTs to generate a more complex three-dimensional waveform (*Omoto et al., 1999*). Recent cryo-electron microscopy studies have elucidated the CA structure in *Chlamydomonas reinhardtii*, which consists of microtubule inner proteins, eleven projections (C1a–f and C2a–e), and a bridge connecting C1 and C2 MTs (*Gui et al., 2022; Han et al., 2022*). Among these projections, only three proteins have been localized to the C2a projection: FAP65 (CCDC108 in mammals), FAP147 (MYCBPAP in mammals), and FAP70 (CFAP70 in mammals) (*Gui et al., 2022; Han et al., 2022; Hou et al., 2021*). Based on the predicted *Chlamydomonas* C2a structure, FAP147 forms the C2a stalk, and FAP65 and FAP70 associate with FAP147 to localize at the base and middle regions of C2a, respectively (*Gui et al., 2022*). The C2a projection is predicted to be anchored on the C2 microtubule by the helices originating from the PF20 (SPAG16 in mammals) homodimer at the base. However, a recent cryo-electron tomography study provides different locations of the three C2a proteins in mouse sperm flagella (*Zhu et al., 2025*). Thus, the C2a projection may vary in its architecture, protein composition, and likely function among species. It is essential to determine the protein composition and the roles of the C2a projection in vertebrates.

In humans, mutations in genes encoding the C2a proteins CCDC108, MYCBPAP, and CFAP70 can lead to multiple morphological abnormalities of the sperm flagella (MMAF), resulting in male infertility (*Beurois et al., 2019; Li et al., 2020; Wang et al., 2021; Wang et al., 2019; Zhou et al., 2025*). However, it remains unclear whether and how the disruption of C2a proteins affects motile cilia in epithelial multiciliated cells (MCCs). In this study, we employed three knockout (KO) mouse models (*Ccdc108* KO, *Mycbpap* KO, and *Cfap70* KO) to systematically investigate the roles of each C2a component in ciliary motility and their functional relationships. Importantly, we found that the loss of any of the C2 proteins in mice caused severe PCD-related phenotypes. CCDC108, MYCBPAP, and CFAP70 interact with each other to form a complex interaction network that is essential for their ciliary localization in motile cilia. Notably, we combined biochemical and microscopic approaches and identified additional C2a components, ARMC3 and MYCBP. Overall, our results reveal the protein composition and physiological functions of the C2a projection and provide significant insights into how this structure contributes to PCD pathology.

## Results

### Generation of KO mouse models for *Ccdc108*, *Mycbpap*, and *Cfap70* using CRISPR/Cas9

Although many CA proteins have been identified by recent proteomic analyses using *Chlamydomonas* (*Dai et al., 2020; L. Zhao et al., 2019*), only FAP65, FAP147, and FAP70 have been designated as C2a proteins (Figure 1A) (*Hou et al., 2021*). Consistently, we found that their corresponding mouse homologs (*Ccdc108*, *Mycbpap*, and *Cfap70*) exhibited the highest expression level in the testis and moderate levels in tissues abundant with motile cilia, such as the reproductive and respiratory systems (Figure 1—figure supplement 1A–C). To investigate the effects of C2a protein deficiency on motile cilia in vertebrates, three KO mouse strains with mutant alleles of *Ccdc108*, *Mycbpap*, or *Cfap70* were developed using CRISPR/Cas9 (Figure 1B).

**Figure 1.**
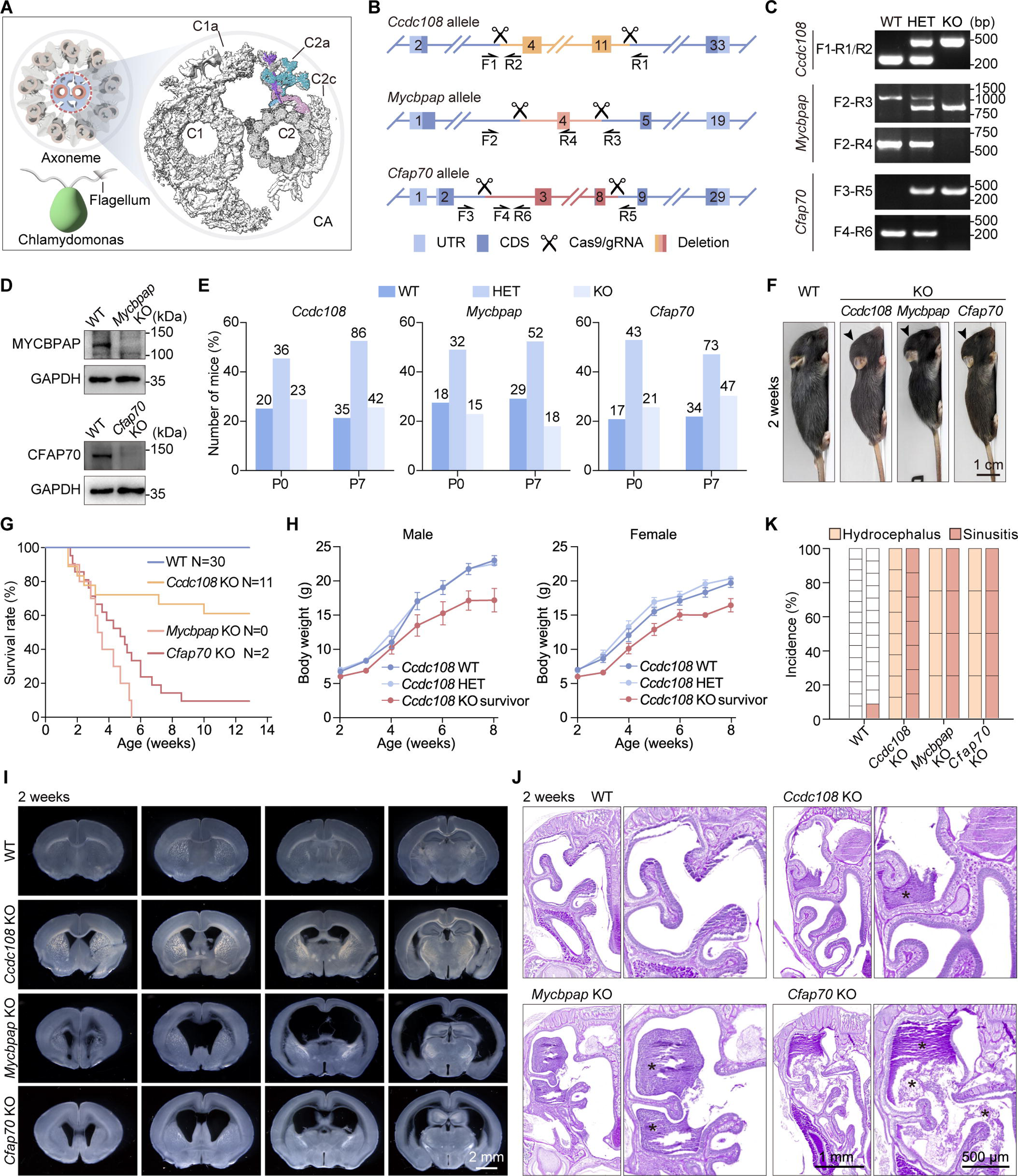
Loss of C2a proteins in mice leads to PCD-related phenotypes. (**A**) Schematics of the cross-section of motile cilia and major projections associated with the C1 and C2 microtubules. The molecular model of the C2a projection (PDB: 7SOM) is superimposed onto the cryo-EM density map from Han, L. et al (*Han et al., 2022*). (**B**) Schematic strategies for the generation of *Ccdc108* KO, *Mycbpap* KO, and *Cfap70* KO mice using CRISPR/Cas9. The genomic positions of the primers used for genotyping are indicated. UTR, untranslated region; CDS, coding sequence. (**C**) Genotyping of WT, HET, and KO mice for each strain. (**D**) Immunoblotting showing the depletion efficiency in *Mycbpap* KO and *Cfap70* KO mEPCs. GAPDH is used as a loading control. (**E**) Genotype distribution profiles of pups at P0 and P7 resulting from matings of HET mice with specified genotypes. Note that *Mycbpap* KO mice were born at Mendelian ratios but experienced early death within one week of birth. (**F**) Representative images of mice with specified genotypes at two weeks of age. (**G**) Survival curves of WT (starting number: 30), *Ccdc108* KO (starting number: 18), *Mycbpap* KO (starting number: 10), and *Cfap70* KO (starting number: 21) mice. The numbers of surviving mice for each genotype at twelve weeks of age are shown. (**H**) Weights of male and female WT, *Ccdc108* HET, and *Ccdc108* KO (survivor) mice were recorded from one to eight weeks of age. Data are presented as mean ± SD (nL=L6 mice per genotype). (**I**) Representative images of serial vibratome sections of the brains from mice with specified genotypes at two weeks of age. (**J**) Periodic Acid-Schiff (PAS) staining of the nasal cavities of mice with specified genotypes at two weeks of age. Magnified images are shown on the right. The asterisks indicate mucus accumulation. (**K**) Incidence of hydrocephalus and sinusitis in mice with specified genotypes. Hydrocephalus and sinusitis were determined as described in (**I**) and (**J**), respectively. The cell number in each column indicates the number of mice analyzed.

The mouse *Ccdc108* gene (ENSMUST00000094844.4) contains 33 exons and encodes a protein of 1,847 amino acids (aa). To generate the *Ccdc108* KO mouse model, a pair of guide RNAs (gRNAs) targeting introns 3 and 11 was designed to deplete exons 4-11, which contain 1,681 base pairs (bp) of coding sequence, causing a frameshift and introducing a premature stop codon in exon 12 (Figure 1B). The generation of the *Mycbpap* KO mouse strain was achieved by depleting exon 4 of the mouse *Mycbpap* transcript (ENSMUST00000093945.10), which consists of 19 exons and encodes a protein of 931 aa. The removal of exon 4 results in a 104-bp loss in the coding region and introduces a premature stop codon in exon 5 (Figure 1B). The mouse *Cfap70* (ENSMUST00000056073.14) encodes a protein of 1,141 aa. A frameshift mutation was created by introducing an 824-bp deletion of exons 3-8 in the mouse *Cfap70* (ENSMUST00000056073.14) (Figure 1B).

The heterozygous (HET) animals from each strain were viable and indistinguishable from their wild-type (WT) littermates. Subsequent matings between HET mice within each strain were used to produce homozygous KO mice. Polymerase chain reaction (PCR) and immunoblotting were performed to confirm genome editing and protein loss in the KO mice, respectively (Figure 1C, D). Since the anti-CCDC108 antibody did not work well for immunoblotting, the loss of CCDC108 in *Ccdc108* KO mice was confirmed by immunostaining of isolated mouse tracheal MCCs (Figure 1—figure supplement 1D). Collectively, these results indicate the successful establishment of KO mouse models for *Ccdc108*, *Mycbpap*, and *Cfap70*.

### Mice lacking CCDC108, MYCBPAP, or CFAP70 exhibit phenotypes related to PCD

Genotyping at postnatal day 0 (P0) revealed that *Ccdc108* KO pups, *Mycbpap* KO pups, and *Cfap70* KO pups were all born at the expected Mendelian ratios; however, the ratio of *Mycbpap* KO mice at P7 appeared to decrease (Figure 1E), indicating that the loss of MYCBPAP may lead to early death. Notably, *Ccdc108* KO mice, *Mycbpap* KO mice, and *Cfap70* KO mice grew more slowly and appeared smaller than WT mice at P14 (Figure 1F). About 10% of the *Mycbpap* KO mice and 5% of the *Cfap70* KO mice died each week starting from one week of age, and nearly all *Mycbpap* KO mice perished before reaching sexual maturity. In contrast, *Ccdc108* KO mice died at a lower rate and in smaller numbers, with approximately 61% surviving by twelve weeks after birth (Figure 1G). Monitoring the body weight of pups from P7 to P56 revealed a significant decrease in weight for both male and female *Ccdc108* KO survivors (Figure 1H), aligning with the smaller body size observed in *Ccdc108* KO mice (Figure 1F). Moreover, *Mycbpap* KO mice and *Cfap70* KO mice all developed dome-shaped heads around two weeks after birth, but this was less common in *Ccdc108* KO mice of the same age (Figure 1F).

Since a dome-shaped head is frequently linked to hydrocephalus, we examined the brain sections of these KO mice at two weeks of age. As shown in Figure 1I, compared to WT mice, *Mycbpap* KO mice and *Cfap70* KO mice both exhibited extensively dilated brain ventricles with a thinner cerebral cortex. However, the enlargement of the brain ventricles in two-week-old *Ccdc108* KO mice appeared less pronounced (Figure 1I). We therefore inferred that hydrocephalus might accelerate mortality in these KO mice. Consistently, when examining the *Ccdc108* KO survivors at eight weeks old, we also observed more dilated ventricles, although most of their heads appeared normal (Figure 1—figure supplement 1E, F). Moreover, histological analysis of the paranasal cavities showed abundant accumulation of protein-rich mucus in KO mice of each strain (Figure 1J), indicating defective mucociliary clearance and sinusitis. Importantly, the hydrocephalus and sinusitis phenotypes were fully penetrant in the KO mice of each strain (Figure 1K). Together, these results showed that *Ccdc108* KO mice, *Mycbpap* KO mice, and *Cfap70* KO mice all developed PCD-related phenotypes.

### Motile cilia in *Ccdc108* KO, *Mycbpap* KO, and *Cfap70* KO mice exhibit abnormal rotational motion due to axonemal ultrastructure defects

Considering these phenotypes in KO mice are associated with defective motile cilia, we next examined motile cilia in the brain ependyma and the upper airway. Scanning electron microscopy (SEM) was carried out to image cilia bundles in the ventricular ependyma. As shown in Figure 2—figure supplement 1A, compared to WT samples, each ciliary bundle in the ependyma of all three KO strains appeared to have fewer motile cilia. Mouse glial cells from the subventricular zone of newborn mice can be cultured to differentiate into multiciliated mouse ependymal cells (mEPCs) (*H. Zhao et al., 2019*). To investigate the impact of the C2a protein loss on multiciliogenesis, we examined motile cilia in mEPC cultures derived from WT and KO brains through immunostaining with acetylated α-tubulin and CEP164 as markers for cilia and centrioles, respectively. Consistently, three-dimensional structured illumination microscopy (3D-SIM) results revealed a reduction in both cilia and centrioles in *Ccdc108* KO, *Mycbpap* KO, and *Cfap70* KO mEPCs (Figure 2A, B), although the decrease in motile cilia was minor (cilia number: 48 ± 22 in WT; 39 ± 14 in *Ccdc108* KO; 31 ± 14 in *Mycbpap* KO; 34 ± 16 in *Cfap70* KO).

**Figure 2.**
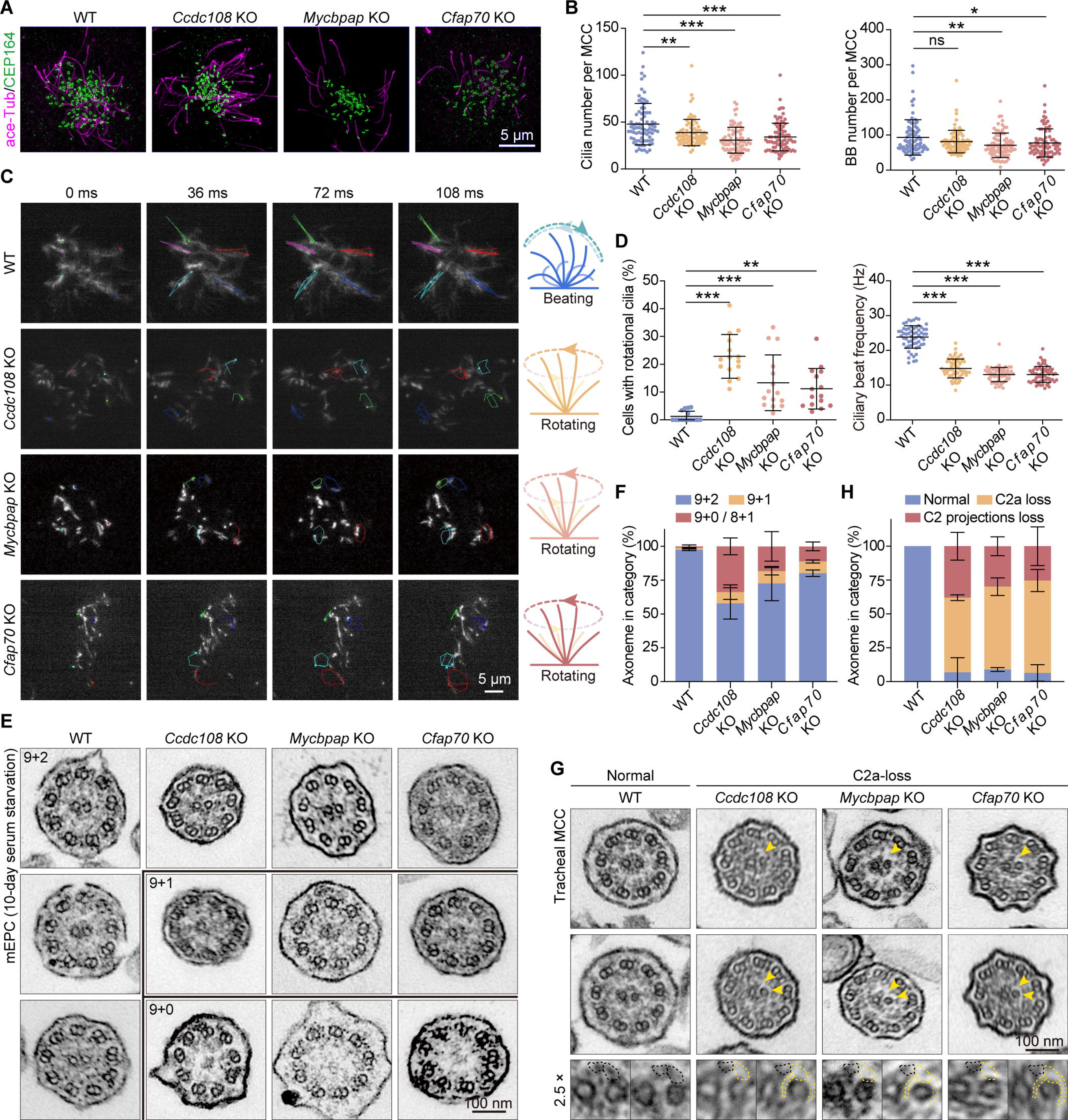
C2a proteins are essential for the integrity of C2-related projections. (**A**, **B**) Immunofluorescence (**A**) and quantifications (**B**) of the number of basal bodies and cilia per mEPC cultured from WT, *Ccdc108* KO, *Mycbpap* KO, and *Cfap70* KO mice. Cells were immunostained with acetylated α-tubulin (ace-Tub) and CEP164 antibodies, and imaged with 3D-SIM. 80 cells from 3 mice (per genotype) were scored using ImageJ. (**C**, **D**) Representative frames (**C**) and quantifications (**D**) of indicated movement modes and beat frequencies of motile cilia in WT, *Ccdc108* KO, *Mycbpap* KO, and *Cfap70* KO mEPCs. Trajectories of four or five cilia in each cell are shown. mEPCs in which the majority of motile cilia displayed rotational movement were considered ‘cells with rotational cilia’. Diagrams illustrate the corresponding ciliary beat patterns. 60 cells from 3 mice (per genotype) were scored using ImageJ. (**E**–**H**) TEM images and quantifications of ciliary axonemes in mEPCs serum-starved for 10 days (**E**, **F**) and in tracheal MCCs (**G**, **H**) from WT, *Ccdc108* KO, *Mycbpap* KO, and *Cfap70* KO mice. Arrowheads indicate the positions of the C2 projections. At least 50 axonemes from 3 mice (per genotype) were scored. Data in (**B**, **D**, **F**, **H**) are presented as mean ± SD. One-way ANOVA with a Dunnett’s test was performed. *, P < 0.05; **, P < 0.01; ***, P < 0.001; ns, not significant.

Given the essential role of the CA in regulating ciliary motility (*Lyu et al., 2024*), we performed high-speed video microscopy to visualize ciliary motility in mEPCs. We noticed that portions of motile cilia in *Ccdc108* KO, *Mycbpap* KO, and *Cfap70* KO mEPCs displayed rotational movement, while control WT cilia beat in an orderly manner with a planar pattern from the top views (Figure 2C, D and Figure 2—figure supplement 2). Moreover, the ciliary beat frequency (CBF) in these KO mEPCs was significantly lower than that of WT motile cilia (Figure 2D and Figure 2—figure supplement 2). Transmission electron microscopy (TEM) was subsequently conducted to analyze the axoneme ultrastructure in mEPCs. We found that motile cilia in *Ccdc108* KO, *Mycbpap* KO, and *Cfap70* KO mEPCs serum-starved for 10 days exhibited characteristic defects in the CP-MTs, including loss of one or both CP-MTs (‘9 + 1’ or ‘9 + 0’ cilia) and transposition of the outer DMT to the center of the axoneme (‘8 + 1’ cilia) (Figure 2E, F). In addition, we examined the motile cilia in the tracheal MCCs of WT and KO mice using SEM and TEM. SEM results revealed no apparent morphological differences between samples of WT and the three KO strains (Figure 2—figure supplement 1B). Strikingly, although no CP-MT defects were observed in the tracheal motile cilia, almost all the examined axonemes showed the loss of C2a or even the loss of entire C2 projections (Figure 2G, H). These results indicate that C2a proteins may play a role in CP-MT maintenance or mechanical stabilization. To test this hypothesis, we examined the axoneme ultrastructure in *Ccdc108* KO mEPCs that were serum-starved for 5 days. Quantification showed that the percentage of axonemes with defective CA decreased further compared to that in *Ccdc108* KO mEPCs serum-starved for 5 days (Figure 2F and Figure 2—figure supplement 1C, D). Overall, these results demonstrate the crucial roles of CCDC108, MYCBPAP, and CFAP70 in maintaining the integrity of the C2a structure, which is required for axoneme stability and proper CBF.

### CCDC108, MYCBPAP, and CFAP70 localize to the axonemal central lumen with mutual interactions

Next, we performed co-immunoprecipitation (co-IP) to investigate whether and how these C2a proteins interact. Immunoprecipitates were prepared from testis lysates using antibodies against MYCBPAP and CFAP70 and analyzed by immunoblotting. We found that CFAP70 was present in the MYCBPAP immune complex, and vice versa (Figure 3A, B). Due to the lack of a specific antibody for CCDC108, we examined its interactions with MYCBPAP and CFAP70 using exogenously expressed proteins in human HEK293T cells. As shown in Figure 3—figure supplement 1A, HA-tagged CCDC108 could be co-immunoprecipitated by both MYCBPAP and CFAP70. To corroborate these results, we performed IP experiments in HEK293T cells co-expressing HA-tagged CCDC108 or CFAP70 with GFP-MYCBPAP. GFP-MYCBPAP was detected in both pull-downs with anti-HA antibody (Figure 3C). Similarly, pull-downs with anti-GFP antibody in HEK293T cells expressing GFP-tagged CCDC108 or MYCBPAP with HA-CFAP70 revealed the presence of HA-CFAP70 (Figure 3D).

**Figure 3.**
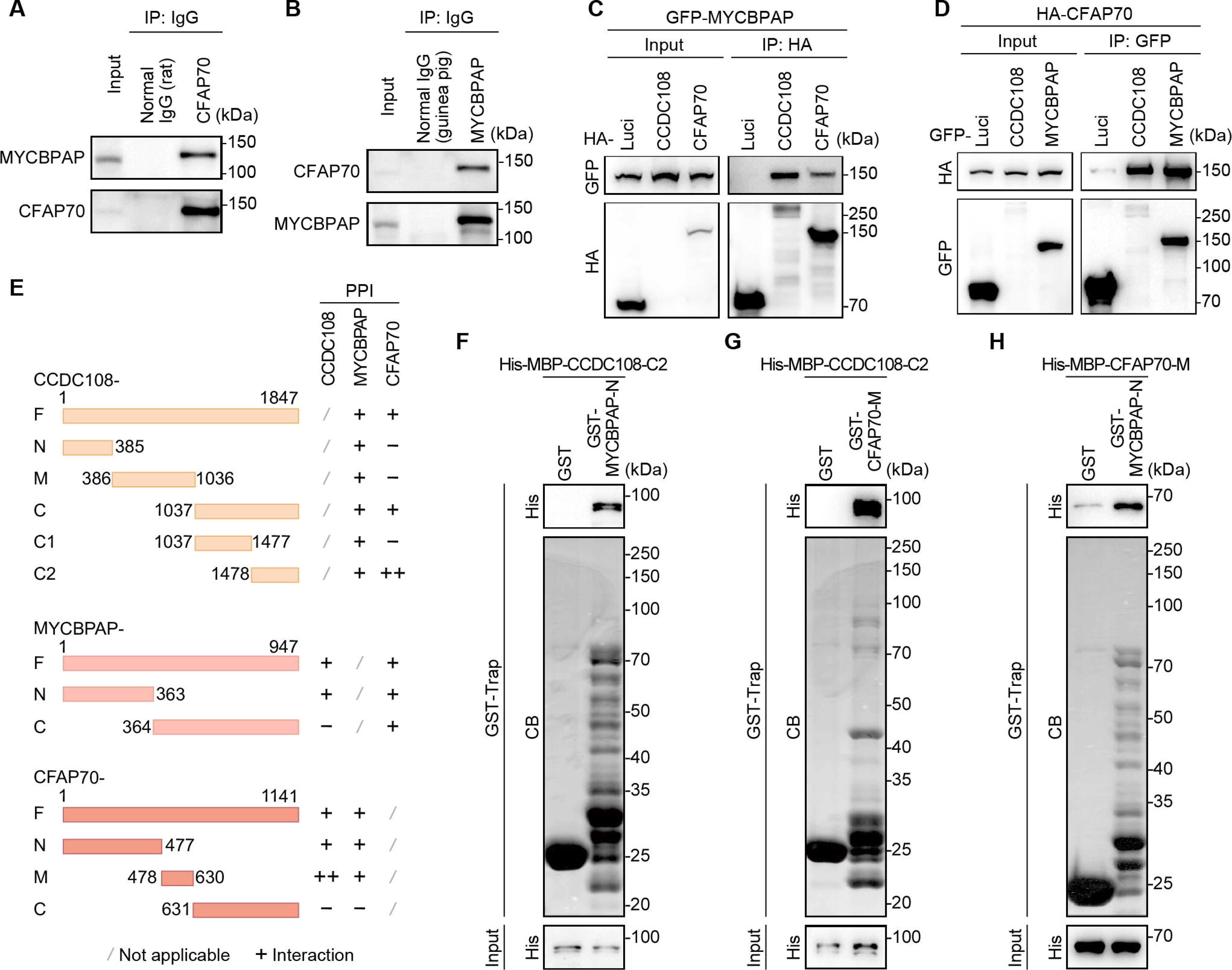
CCDC108, MYCBPAP, and CFAP70 mutually interact with each other. (**A**, **B**) Co-IP and immunoblotting showing the interaction between endogenous MYCBPAP and CFAP70. Co-IP was performed with a normal rat IgG control antibody and a rat polyclonal anti-CFAP70 antibody (**A**) or a normal guinea pig IgG and a guinea pig polyclonal anti-MYCBPAP antibody (**B**) in mouse testis lysates. (**C**, **D**) Co-IP and immunoblotting analyses in HEK293T cells exogenously expressing indicated proteins. HA-tagged proteins in (**C**) and GFP-tagged proteins in (**D**) were immunoprecipitated with anti-HA and anti-GFP agarose beads, respectively. Blots were probed with the indicated antibodies. Luci, luciferase. (**E**) Diagrams of CCDC108, MYCBPAP, and CFAP70 full-length (F) and truncated fragments showing their ability to interact. Interactions were determined through co-IP analyses. Numbers indicate amino acid positions. PPI, protein-protein interaction. (**F**–**H**) GST pull-down showing direct interactions using purified fragment proteins. Blots were probed with the indicated antibodies. CB, coomassie blue staining.

To identify the domains responsible for their interactions, we generated a series of truncation mutants for each protein and performed co-IP experiments. We found that all the CCDC108 fragments could bind MYCBPAP, but only the C-terminal region was essential for its association with CFAP70 (Figure 3E and Figure 3—figure supplement 1B, C). Conversely, MYCBPAP interacted with CCDC108 via its N-terminus, while both the N- and C-terminal fragments could bind to CFAP70 (Figure 3E and Figure 3—figure supplement 1D, E). Regarding CFAP70, we observed that the two fragments, CFAP70-N (1-477 aa) and CFAP70-M (478-630 aa), could associate with both CCDC108 and MYCBPAP, whereas its C-terminal fragment lacked the binding capacity (Figure 3E and Figure 3—figure supplement 1F, G). Additionally, GST pull-down experiments with purified recombinant proteins from *Escherichia coli* lysates further confirmed direct interactions among the three C2a proteins (Figure 3F–H). Together, we conclude that the three C2a proteins can directly interact with each other.

Next, we examined the cellular localization of C2a proteins in mEPCs by detecting either GFP-tagged or endogenous proteins. In multiciliated mEPCs expressing GFP-tagged C2a proteins, GFP signal was observed to localize to motile cilia (Figure 4A). 3D-SIM imaging further revealed that GFP signals were located within the axonemal central lumen, which was defined by the axonemal microtubules labeled with the anti-acetylated α-tubulin antibody (Figure 4B). Consistently, endogenous CCDC108, MYCBPAP, and CFAP70 were also detected within the axonemal central lumen (Figure 4C). Additionally, MYCBPAP was observed to co-localize with CCDC108 in the ciliary central lumen and with CFAP70 (Figure 4D). Overall, our results demonstrate that the three C2a proteins, CCDC108, MYCBPAP, and CFAP70, colocalize in the central lumen of motile cilia and interact with each other.

**Figure 4.**
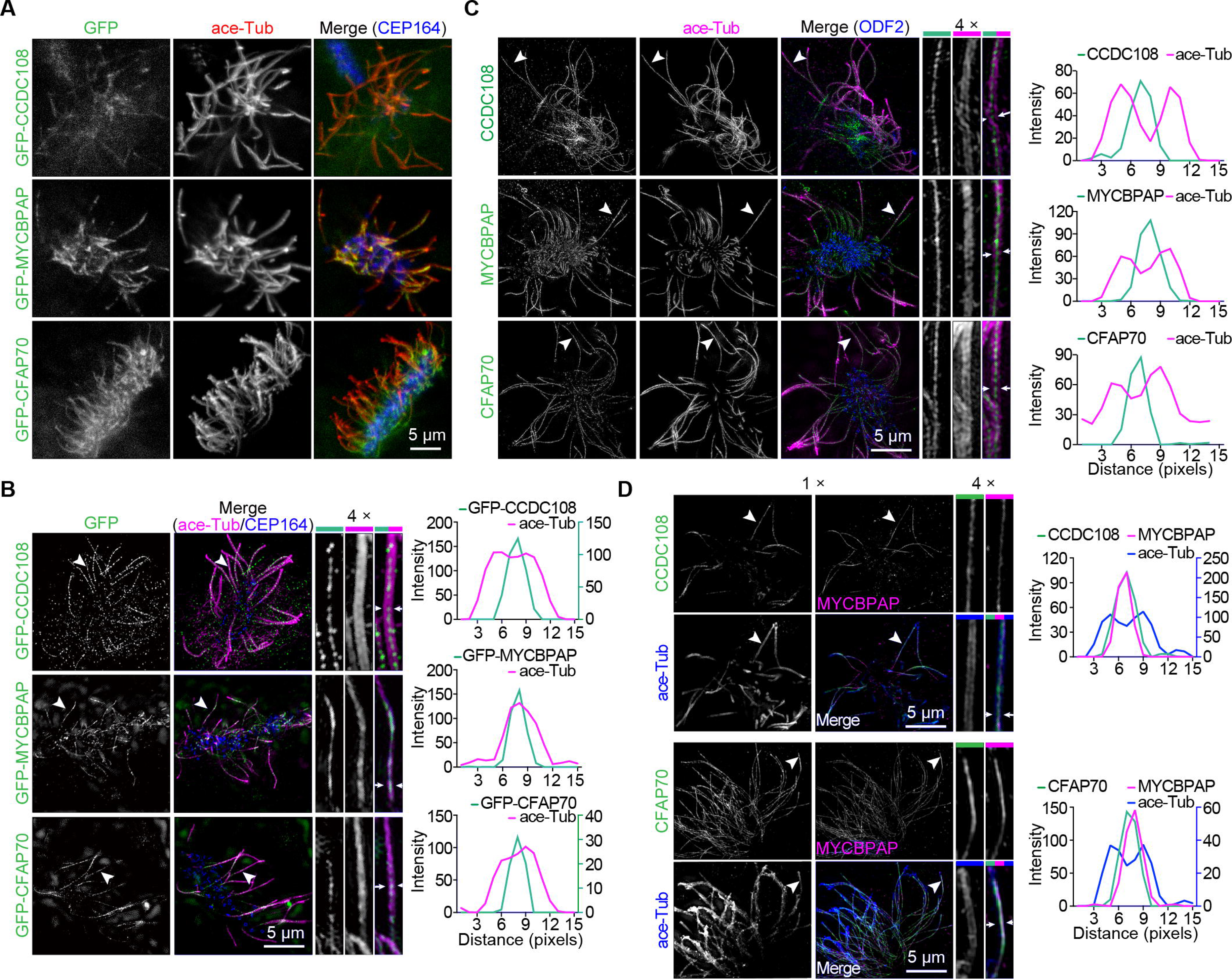
CCDC108, MYCBPAP, and CFAP70 localize to the axonemal central lumen. (**A**, **B**) Confocal (**A**) and 3D-SIM (**B**) images of mEPCs expressing GFP-tagged proteins immunostained with the indicated antibodies. Magnified images of the cilia indicated by arrowheads are shown on the right. Line-scan graphs show the immunofluorescence intensity along the positions marked by arrows in the magnified images. The right Y-axis describes the immunofluorescence intensity of GFP-CCDC108 and GFP-CFAP70, respectively. (**C**, **D**) 3D-SIM images of mEPCs immunostained with the indicated antibodies. Magnified images of the cilia indicated by arrowheads are shown on the right. Line-scan graphs show the immunofluorescence intensity along the positions marked by the two arrows in the magnified images. The right Y-axis describes the immunofluorescence intensity of ace-Tub in (**D**).

### CCDC108, MYCBPAP, and CFAP70 are all essential for the stable docking of each C2a protein

Given the mutual interactions and ciliary colocalization of the C2a components, we set out to determine the localization dependencies among the three C2a proteins. mEPCs from *Ccdc108* KO, *Mycbpap* KO, and *Cfap70* KO brains were cultured ex vivo and subjected to immunostaining. In control WT cells, CCDC108 staining was located in motile cilia (Figure 5A). In contrast, in both *Mycbpap* KO and *Cfap70* KO mEPCs, the ciliary CCDC108 signal was barely detectable (Figure 5A). Strikingly, we found that the loss of any of the three C2a proteins could remarkably disrupt the ciliary localization of the other two C2a proteins (Figure 5B, C). A similar phenomenon was also observed in mouse tracheal motile cilia from *Ccdc108* KO, *Mycbpap* KO, and *Cfap70* KO mice (Figure 5D–F). However, the ciliary localization of SPEF1 (a CP-MT-seam binding protein (*Legal et al., 2025; Zheng et al., 2019*)), HYDIN (an essential component of the C2b projection (*Lechtreck et al., 2008; Lechtreck and Witman, 2007*)), and RSPH3 (a core radial spoke protein (*Gui et al., 2021; Jeanson et al., 2015; Meng et al., 2024*)) was not apparently affected (Figure 5—figure supplement 1A–C).

**Figure 5.**
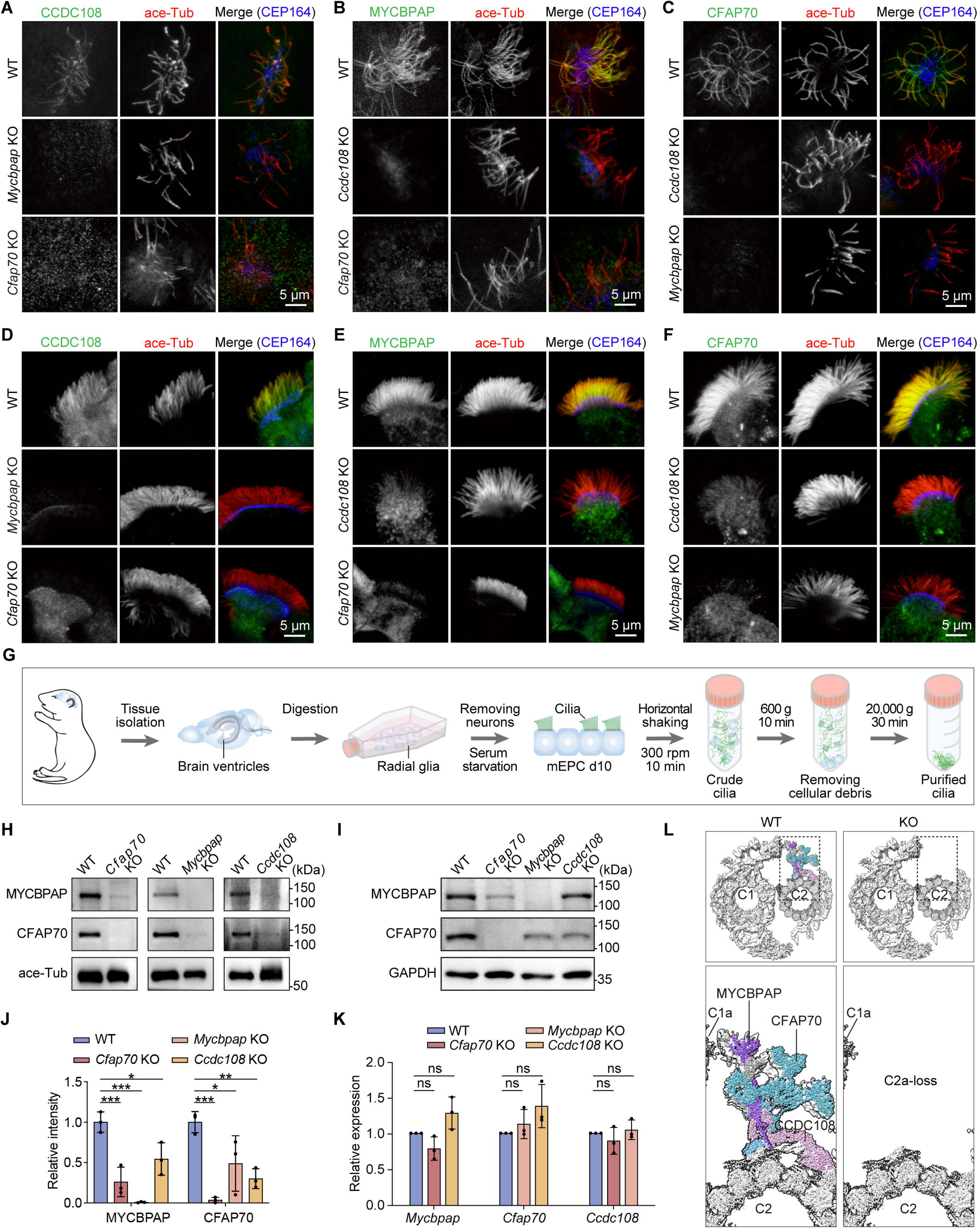
CCDC108, MYCBPAP, and CFAP70 are all essential for the stable docking of each C2a protein. (**A**–**F**) Representative confocal images of mEPC (**A**–**C**) and tracheal MCCs (**D**–**F**) from WT, *Ccdc108* KO, *Mycbpap* KO, or *Cfap70* KO mice immunostained with the indicated antibodies. Note that the ciliary staining of C2a proteins was greatly reduced in KO samples. (**G**) Schematic diagrams of mEPC culture and motile cilia purification. (**H**–**J**) Immunoblotting and quantification showing the C2a protein levels in motile cilia (**H**) or mEPCs (**I**) derived from *Ccdc108* KO, *Mycbpap* KO, and *Cfap70* KO mEPCs. Acetylated α-tubulin (ace-Tub) and GAPDH are used as loading controls, respectively. The MYCBPAP or CFAP70 band intensity in (**I**) was normalized to the corresponding GAPDH intensity. Data are presented as mean ± SD. (**K**) Real-time PCR analyses showing the expression levels of the indicated genes in *Ccdc108* KO, *Mycbpap* KO, and *Cfap70* KO mEPCs. The expression was normalized using the corresponding *Gapdh* as the reference gene and WT as the reference sample (△△C_T_ method). Data are from three independent biological repeats and are presented as mean ± SD. (**L**) Proposed model showing the absence of the C2a projection in *Ccdc108* KO, *Mycbpap* KO, and *Cfap70* KO axoneme. One-way ANOVA with a Dunnett’s test was performed. *, P < 0.05; **, P < 0.01; ***, P < 0.001; ns, not significant.

In addition, we purified motile cilia from WT and KO mEPCs (*Hao et al., 2022*) and examined the ciliary protein levels of C2a proteins (Figure 5G). Consistently, ciliary CFAP70 was nearly undetectable in *Mycbpap* KO motile cilia, and the level of ciliary MYCBPAP in *Cfap70* KO cilia was also greatly decreased (Figure 5H). The absence or decreased levels of C2a proteins in motile cilia of *Ccdc108* KO, *Mycbpap* KO, and *Cfap70* KO mEPCs may result from either a reduction in the overall amount of each C2a protein or an inability to form a stable C2a projection in the cilia. To investigate this, we measured the protein levels of C2a proteins in *Ccdc108* KO, *Mycbpap* KO, and *Cfap70* KO mEPCs. Immunoblotting of whole cell lysates revealed a dramatic decrease in MYCBPAP and CFAP70 levels in *Ccdc108* KO, *Mycbpap* KO, or *Cfap70* KO mEPCs (Figure 5I, J). However, quantitative PCR showed no significant difference in mRNA levels between WT and KO mEPCs (Figure 5K), suggesting that losing any of the three C2a proteins may make other components of the C2a projection more vulnerable to proteolytic degradation. Given the ultrastructural defects in KO motile cilia observed by TEM (Figure 2G), our results demonstrate the essential role of each C2a protein in the stable docking of other C2a components, thereby maintaining the integrity of the C2a projection (Figure 5L).

### Identification of ARMC3 and MYCBP as new C2a components

We have thus far characterized CCDC108, MYCBPAP, and CFAP70 as essential C2a components in mouse motile cilia. Next, we examined whether additional proteins are present in this projection. To identify new components of the C2a structure, we employed affinity chromatography and mass spectrometry (MS) to detect proteins that interact with endogenous MYCBPAP and CFAP70 in lysates from mouse testes (Figure 6A–D). MS analysis detected peptides from known binding proteins for MYCBPAP, such as CCDC108, and CFAP70, as well as peptides for known CFAP70 binding proteins, including CCDC108 and MYCBPAP (Figure 6C, D). Interestingly, ARMC3 and MYCBP were identified in both MS analyses (Figure 6C, D). We then performed co-IP experiments and examined the co-immunoprecipitated proteins using specific antibodies. Immunoblotting showed that endogenous ARMC3 and MYCBP were readily detected in complexes immunoprecipitated with MYCBPAP and CFAP70 from mouse testis lysates (Figure 6E, F). To determine whether these interactions are conserved across vertebrate motile ciliated cells, we performed co-immunoprecipitation using lysates from mouse trachea and cultured mEPCs. We found that, in both tracheal and mEPC lysates, CFAP70, ARMC3, and MYCBP were co-immunoprecipitated with MYCBPAP (Figure 6—figure supplement 1A, B), indicating that the interactions among the C2a components are conserved in vertebrate motile ciliated cells. To further confirm these interactions, HA-tagged ARMC3 or MYCBP was co-expressed with GFP-tagged C2a proteins in HEK293T cells and immunoprecipitated with anti-HA agarose beads. As shown in Figure 6G, CCDC108, MYCBPAP, and CFAP70 were all detected in the ARMC3 immunoprecipitated complex. However, only MYCBPAP was present in the MYCBP immunocomplex (Figure 6H). These findings suggest that ARMC3 is likely a constitutive C2a component, whereas MYCBP may associate with specific regions of this structure.

**Figure 6.**
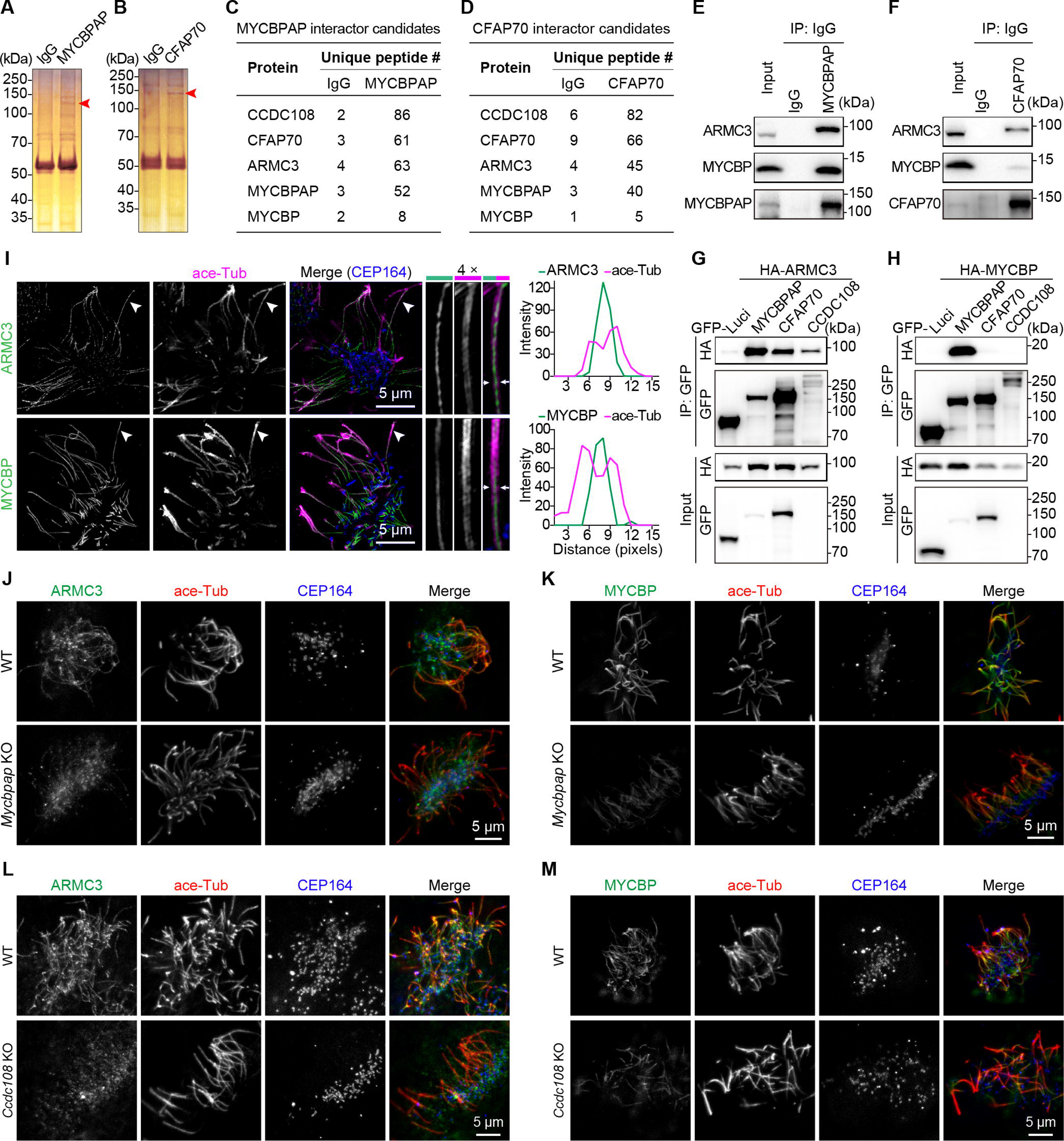
Identification of ARMC3 and MYCBP as new C2a components. (**A**, **B**) Silver staining of proteins immunoprecipitated from mouse testis lysates using a normal guinea pig IgG and a guinea pig polyclonal anti-MYCBPAP antibody (**A**) or a normal rat IgG control antibody and a rat polyclonal anti-CFAP70 antibody (**B**). The bands of MYCBPAP and CFAP70 are indicated by red arrowheads. (**C**, **D**) Interactor candidates of MYCBPAP (**C**) and CFAP70 (**D**) identified by mass spectrometry analysis. (**E**, **F**) Co-IP and immunoblotting showing the interactions of endogenous MYCBPAP (**E**) and CFAP70 (**F**) with ARMC3 and MYCBP. Co-IP was performed with a normal guinea pig IgG and a guinea pig polyclonal anti-MYCBPAP antibody (**E**) or a normal rat IgG control antibody and a rat polyclonal anti-CFAP70 antibody (**F**) in mouse testis lysates. (**G**, **H**) Co-IP and immunoblotting analyses in HEK293T cells exogenously expressing indicated proteins. GFP-tagged proteins were immunoprecipitated with anti-GFP agarose beads. Blots were probed with the indicated antibodies. Luci, luciferase. (**I**) 3D-SIM images of mEPCs immunostained with the indicated antibodies. Magnified images of the cilia indicated by arrowheads are shown on the right. Line-scan graphs show the immunofluorescence intensity along the positions marked by two arrows in the magnified images. (**J**–**M**) Representative confocal images of mEPC from WT, *Mycbpap*, and *Cfap70* KO mice immunostained with the indicated antibodies.

Proteins involved in cilia formation generally maintain high and sustained expression levels during the later stages of multiciliogenesis (*Nai et al., 2025; Zhao et al., 2022; Zhao et al., 2013*). We observed that the protein levels of ARMC3 and MYCBP gradually increased as mEPCs differentiated, showing a similar expression pattern to that of the known C2a proteins (Figure 6—figure supplement 1C), which further suggests their close association with motile cilia. Next, we examined the subcellular localization of ARMC3 and MYCBP through immunostaining of multiciliated mEPCs. As expected, both ARMC3 and MYCBP localized in the axonemal central lumen (Figure 6I). To better confirm their connection with the C2a structure, we sought to determine how the loss of C2a proteins affects their ciliary localization. Strikingly, compared to WT cells, in *Mycbpap* or *Cfap70* KO mEPCs, the ARMC3 ciliary signal became barely perceptible, and MYCBP appeared much fainter (Figure 6J–M). Collectively, our results demonstrate that ARMC3 and MYCBP are two additional components of the C2a projection in motile cilia.

## Discussion

In this study, we utilized *Ccdc108*, *Mycbpap*, and *Cfap70* KO mouse models to demonstrate the essential roles of these proteins in maintaining ciliary motility and tissue homeostasis. We found that the ciliary incorporation of CCDC108, MYCBPAP, and CFAP70 is essential for one another’s stability, with the loss of any single component triggering C2a collapse and likely leading to the proteolytic degradation of the remaining partners. Our results suggest that the C2a projection relies on a complex interaction network for stable docking within the axoneme, revealing a strict mutually dependent assembly model for this projection in vertebrates.

We significantly expanded the known vertebrate C2a proteome by identifying ARMC3 and MYCBP as new components. ARMC3 interacts with all known C2a proteins and depends on them for ciliary localization, suggesting it is a constitutive component of the projection. Additionally, we identified MYCBP (the homolog of *Chlamydomonas* FAP174 (*Hou et al., 2021; Rao et al., 2016*)) as a new CA protein, which localizes to the axonemal central lumen in vertebrate motile cilia. Interestingly, while MYCBP levels were reduced in C2a KO cilia, the protein was still detectable, supporting the hypothesis that MYCBP may have additional axonemal docking sites beyond the C2a projection, similar to findings in protozoa (*Joachimiak et al., 2021; Rao et al., 2016; L. Zhao et al., 2019*). However, the physiological roles of MYCBP and ARMC3 in vertebrate motile cilia still need further investigation.

Importantly, we find that in C2a-loss mEPCs, most cells exhibit wave-like motion with a significant reduction in CBF, and approximately 25% switch to rotational movement, which aligns with the observation that a similar percentage of KO mEPCs have the CP-MT-loss defect. These results suggest that the C2a projection specifically regulates the ciliary beat cycle. However, the loss of the C2a projection destabilizes other C2-associated projections, ultimately leading to destabilization of the CP-MTs. Interestingly, we observed a striking tissue-specific difference in resilience: while ependymal motile cilia frequently exhibited severe defects such as the loss of one or both CP-MTs, tracheal cilia typically retained the CP-MTs despite losing the C2a projection. Given that interactions among C2a components are conserved across multiple motile ciliated cells, including tracheal and ependymal multiciliated cells, our finding suggests that tracheal cilia may possess distinct molecular or structural properties that render them more resilient to C2a loss than their ependymal counterparts.

Our knockout mouse exhibited fully penetrant hydrocephalus and sinusitis, confirming that defects in C2a proteins drive PCD pathology. In humans, mutations in CCDC108, MYCBPAP, or CFAP70 have been found to cause male infertility due to multiple morphological abnormalities of the sperm flagella (MMAF) (*Beurois et al., 2019; Chen et al., 2023; Li et al., 2020; Lu et al., 2023; Wang et al., 2021; Wang et al., 2019; X. Zhang et al., 2019; Zhou et al., 2025*). However, it has not been fully examined whether these patients developed PCD symptoms. Notably, our models in a C57BL/6 background displayed high mortality and severe phenotypes, whereas models on hybrid backgrounds survived to sexual maturity (*Chen et al., 2023; Wang et al., 2024*). This discrepancy highlights a profound genetic background effect on disease severity, offering a potential explanation for why human PCD patients often present with milder symptoms or exhibit variable penetrance.

In summary, we systematically investigate the functional roles of individual C2a proteins in mouse motile cilia and demonstrate the requirement of each C2a protein for maintaining the integrity of the C2a projection. Importantly, we identify ARMC3 and MYCBP as new components of the C2a projection. These findings reveal the physiological functions of the C2a proteins and provide novel insights into the molecular mechanisms underlying PCD.

## Materials and methods

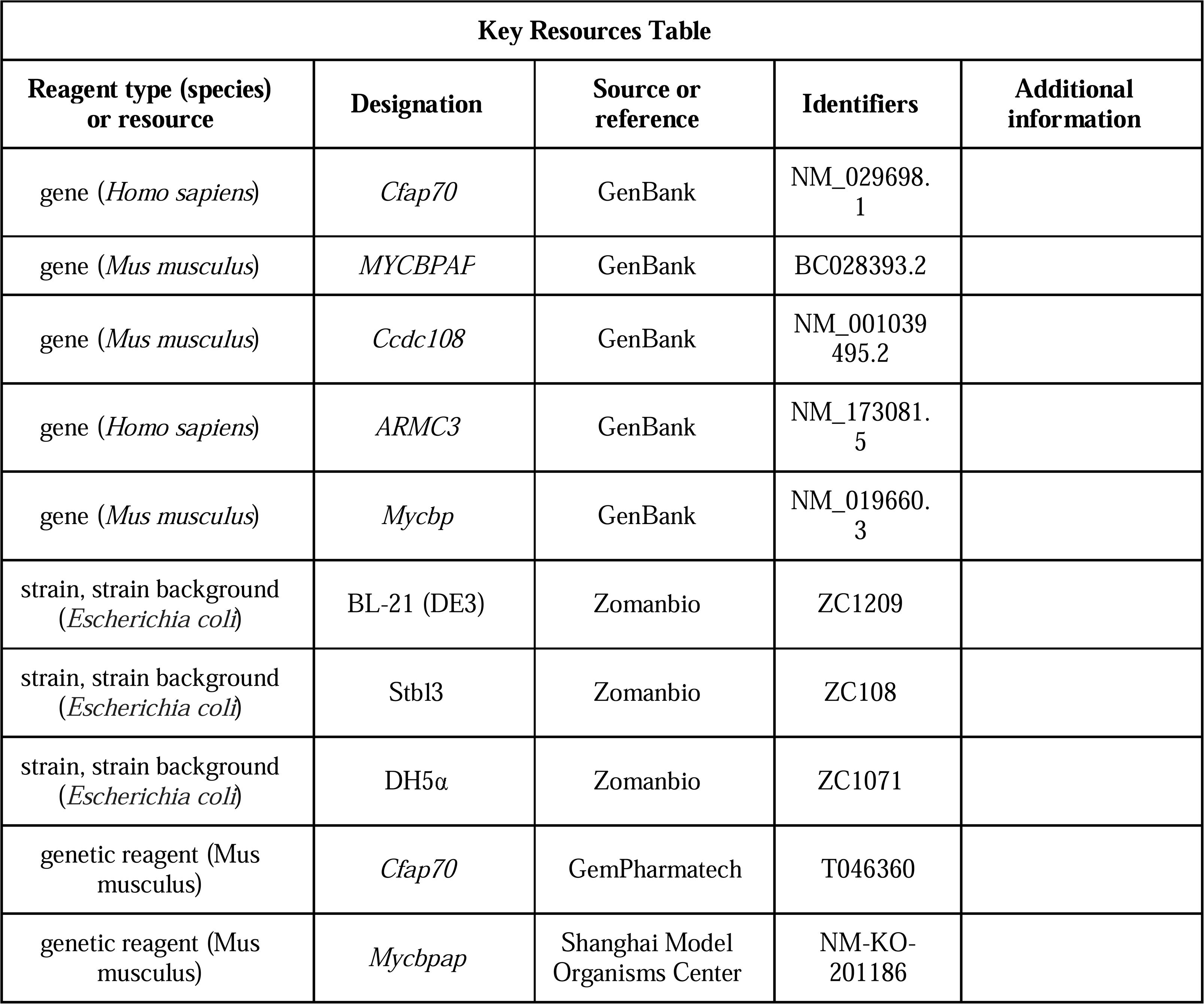
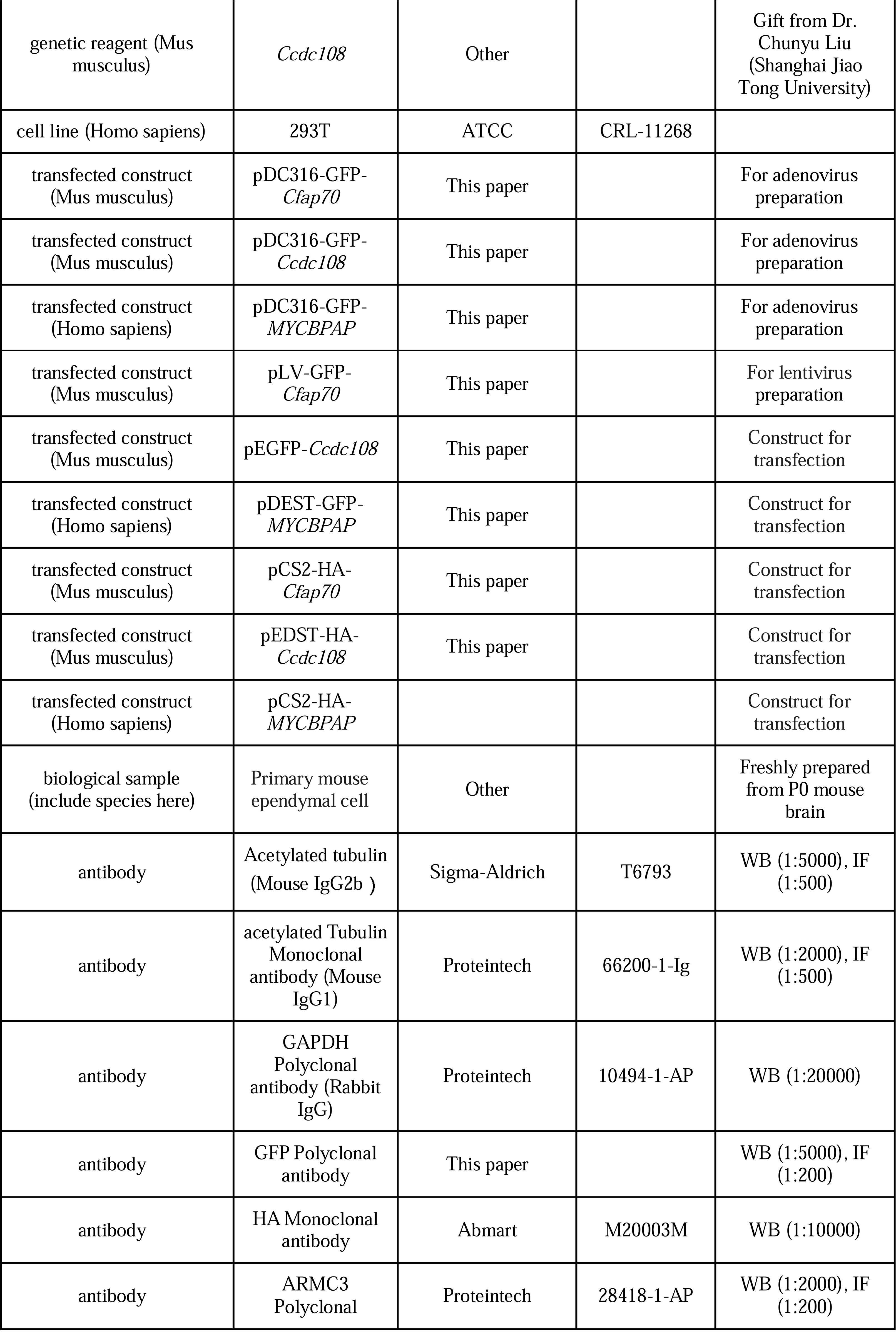
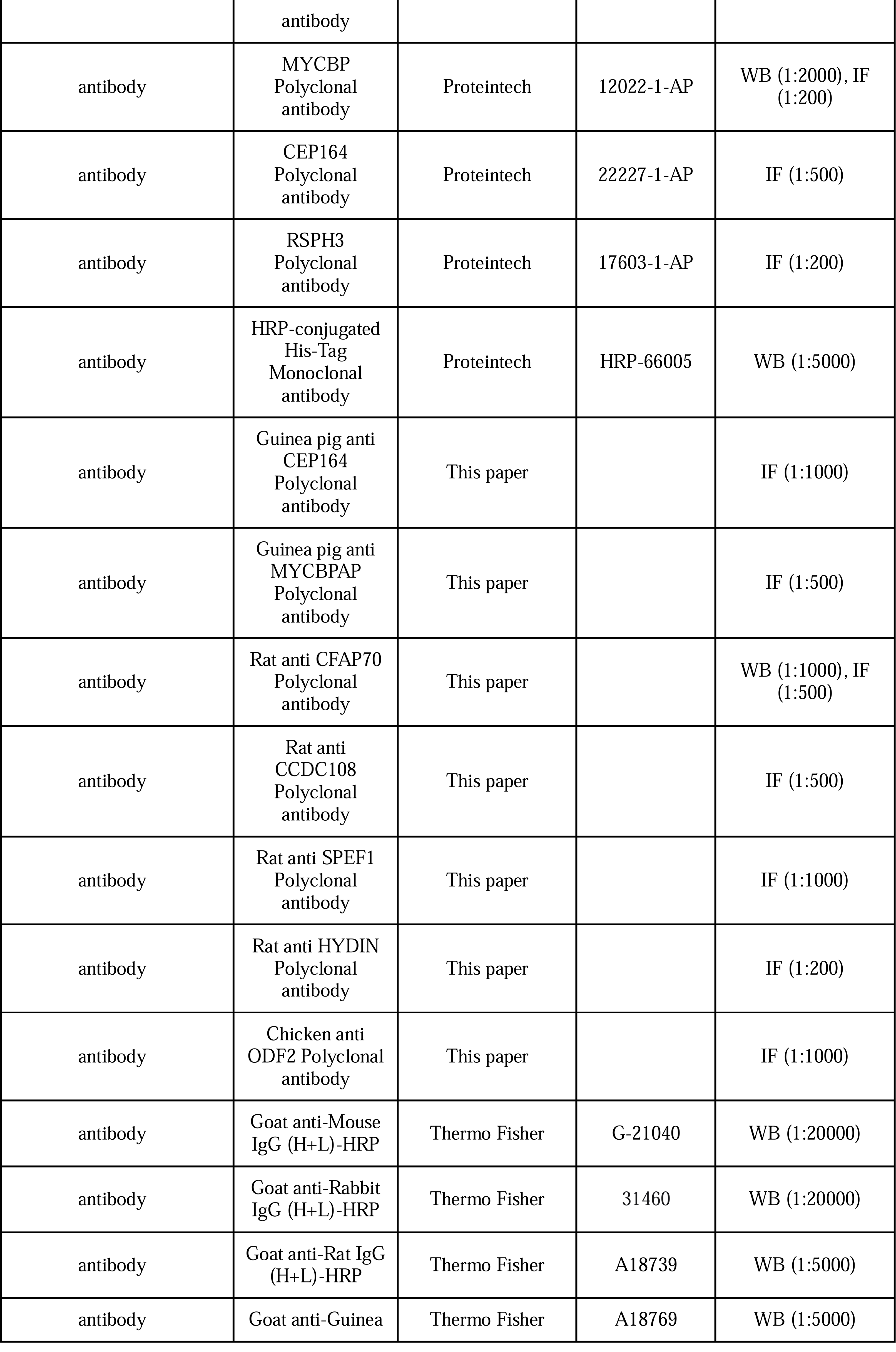
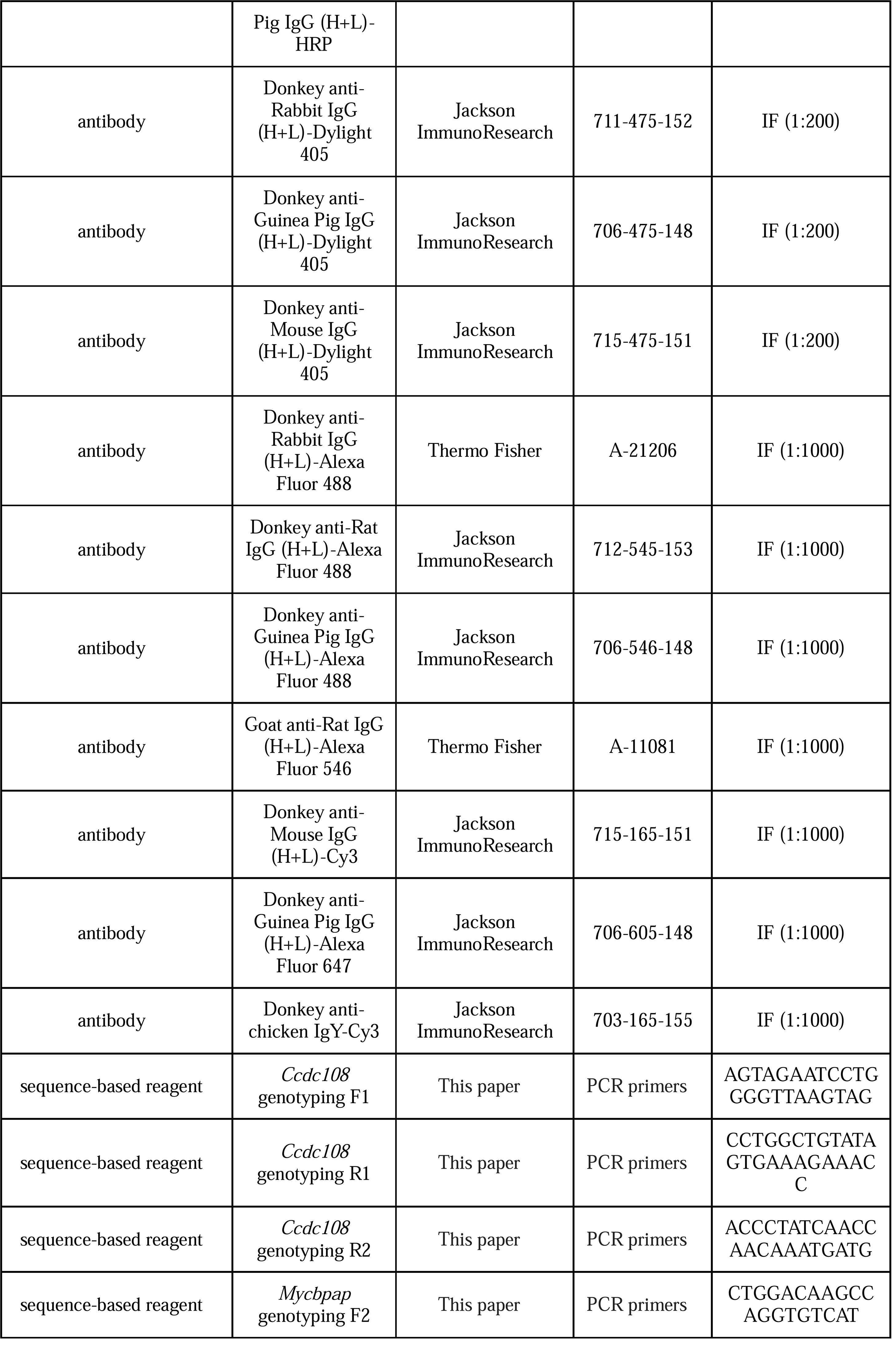
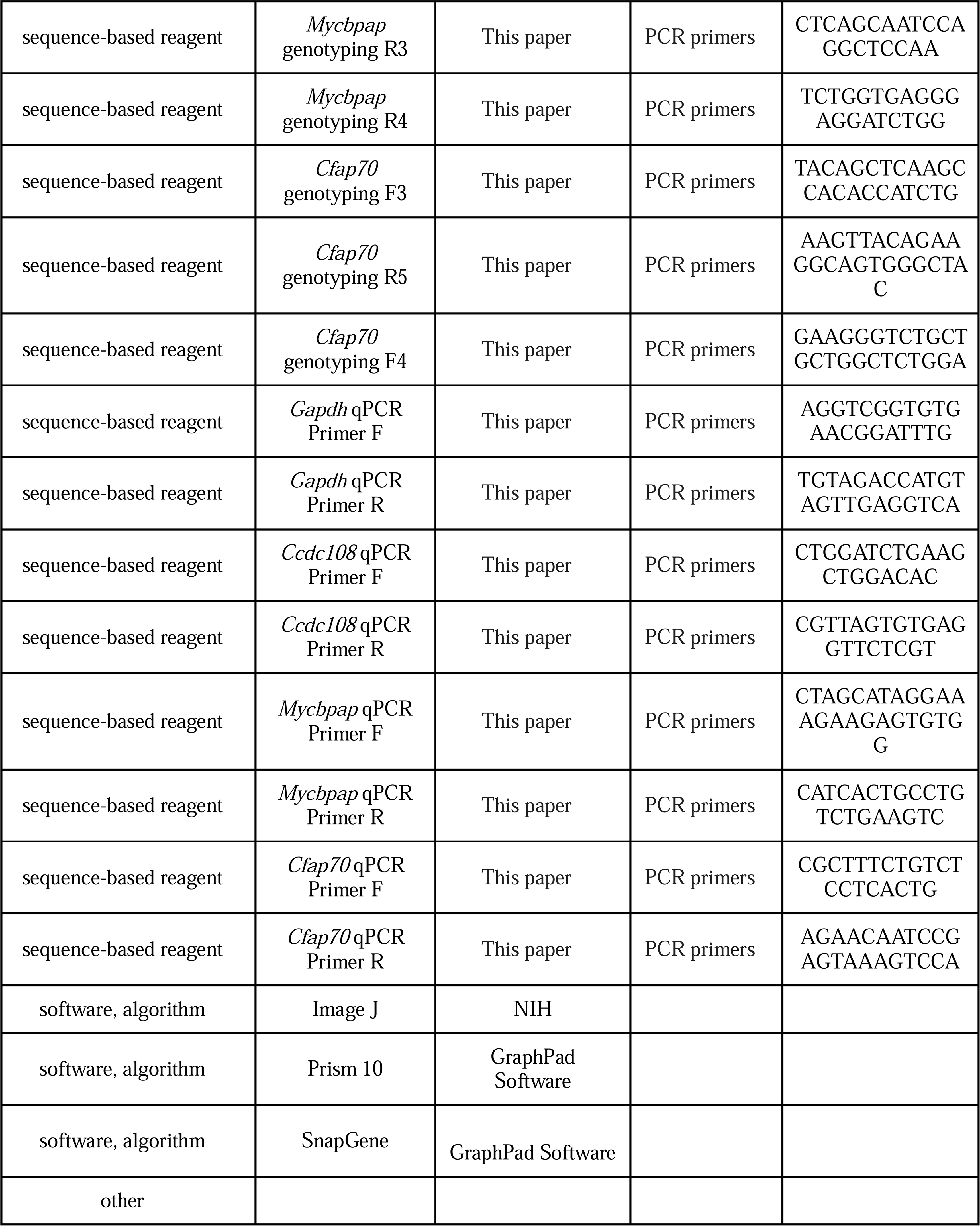

### Mice

All mouse experiments were conducted in accordance with the ethical guidelines of Shandong Normal University and were approved by the Institutional Animal Care and Use Committee (AEECSDNU2022047). The *Ccdc108* KO mouse model was a gift from Dr. Chunyu Liu (Shanghai Jiao Tong University School of Medicine). The *Mycbpap* KO mouse model (NM-KO-201186) and *Cfap70* KO mouse model (T046360) were generated on the C57BL/6J background by the Shanghai Model Organisms Center and GemPharmatech, respectively. Genotyping was performed using 2 × Taq Plus Master Mix II (P213, Vazyme). Fertility test and gene expression patterns were performed as previously described (*Lu et al., 2025; Zi et al., 2024*). qPCR analyses were conducted using the Hieff UNICON advanced qPCR SYBR Master Mix (11185ES08, Yeasen Biotech). *Gapdh* was used for normalization.

### Histological analysis

Mice were anesthetized with an intraperitoneal injection of 1.25% tribromoethanol (Avertin) at a dose of 250 mg/kg. Then, the mice were transcardially perfused with 50 ml of PBS followed by 50 ml of PBS containing 4% paraformaldehyde (PFA). The brains were then dissected, immediately post-fixed in 4% PFA for 24 h at 4°C, and cut into 250-µm-thick sagittal slices. The brain slices were placed into a glass-bottom dish (D35-20-1.5-N, Cellvis) and imaged with a Leica M205FA fluorescence stereo microscope. To examine the mouse nasal sinus, the tissue was isolated, decalcified until soft in a decalcifying solution (10% HCl, 2% acetic acid, 4% PFA, 2.5 M NaCl), and then dehydrated using an automated tissue processor (HistoCore PEARL, Leica). The samples were embedded in paraffin, sectioned at 10 µm thickness with a rotary microtome, deparaffinized with xylene, rehydrated, and stained with the glycogen PAS staining kit (KGE1103-400, Nanjing KeyGen Biotech) and hematoxylin stain kit (E607318, Sangon Biotech). Final treatments included ethanol dehydration gradients, xylene clearing, and sealing.

### Plasmids

Full-length mouse *Ccdc108* (NM_001039495) and *Cfap70* (NM_029698.1) were amplified from a mouse cDNA library via PCR. Full-length human *MYCBPAP* (BC028393.2) was obtained from the DNASU Plasmid Repository (Arizona State University). Full-length and relative fragments were subcloned into the donor vector pDONR221 using the Gateway BP clonase (11789100, ThermoFisher) to obtain entry plasmids. The indicated expression constructs were generated through LR recombination reactions involving the entry plasmids and Gateway destination vectors (Kits #1000000211 and #1000000107, Addgene) using the Gateway LR clonase (11791100, ThermoFisher). To generate the lentiviral plasmids, PCR products were subcloned into the lentiviral vector, pLV-GFP-C1 (*Zhao et al., 2013*), using the ClonExpress II One Step Cloning Kit (C112, Vazyme). For the bacterial expression constructs, fragments encoding amino acids 1-363 of MYCBPAP and 468-630 of CFAP70 were subcloned into pGEX-4T-1, and the entry plasmid containing the fragment encoding amino acids 1,478-1,847 of CCDC108 was used to recombine with the pDEST-566 vector (#11517, Addgene) for expressing the His-MBP fused protein. All constructs were verified via Sanger sequencing analysis.

### Cell culture, transfection, and virus infection

HEK293T (CRL-11268, ATCC) and HEK293A (R70507, Thermo Fisher) cells were maintained in Dulbecco’s Modified Eagle’s Medium (DMEM; C11995500BT, Thermo Fisher) supplemented with 10% fetal bovine serum (FBS; A5256701, Thermo Fisher), 1% penicillin/streptomycin (P1400, Solarbio), and 2 mM L-alanyl-L-glutamine (G0190, Solarbio). Cells were routinely tested for mycoplasma contamination. HEK293T cells were transfected with plasmids and polyethylenimine (PEI; 23966, Polysciences) at a 2:3 ratio, and harvested 48 h after transfection. To produce lentiviral particles, HEK 293T cells were transfected with the lentiviral plasmid, the packaging vector delta 8.9, and the VSV-G envelope glycoprotein vector at the ratio of 5:3:2 using PEI. The culture medium containing lentiviral particles was collected 48 h post-transfection. Adenovirus and multiciliated mEPCs were prepared as described previously (*Chen et al., 2025; Zhao et al., 2021*). To infect mEPCs with lentivirus, the medium containing lentiviral particles was added to the mEPC culture medium at a 2:1 dilution. mEPCs serum-starved for 5 or 10 days were subjected to analyses.

### High-speed live cell imaging

mEPCs cultured in glass-bottom dishes were serum-starved for 10 days to induce cilia formation. Cilia were then labeled with 100 nM SiR-tubulin (SC002, Spirochrome) in serum-free DMEM medium for 1 h and then imaged in an incubation chamber (37 °C, 5% CO2, and 80% humidity) with an Olympus Xplore SpinSR10 microscope equipped with a UPLAPO OHR 60L×L/1.50 oil objective and an ORCA-Fusion camera (Hamamatsu). The laser power for the SiR-tubulin channel (640 nm) was adjusted to 95% to enable imaging at 12 ms intervals. Ciliary trajectories were visualized using the manual tracking plugin in ImageJ. Four or five traceable cilia per cell were tracked over approximately 1000 ms. mEPCs with most motile cilia displaying rotational motility were considered ‘cells with rotational cilia’. The CBF of each cilium was calculated from the overall time of 10 beating cycles (*Y. Zhang et al., 2019*).

### Motile cilia isolation

To purify motile cilia from mEPC cultures, mEPCs were serum-starved for 10 days in a 75-cm² flask and then washed twice with ice-cold PBS and twice with deciliation buffer (20 mM Pipes, 20LmM CaCl_2_, 250 mM Sucrose, pH 5.5). Then, 9 ml of deciliation buffer supplemented with 0.01% Triton X-100 was added to the flask, followed by horizontal shaking for 10 min at 300 rpm in a 37 °C incubator to fully release the apical motile cilia. The suspension containing the cilia was collected and centrifuged for 10 min at 600L×Lg at 4L°C to remove cell debris. The supernatant was further centrifuged at 20,000L×Lg for 30 min at 4 °C. The pellet was lysed with the SDS sampling buffer (50 mM Tris, 2% SDS, 10% glycerol, pH 6.8) for immunoblotting.

### Fluorescence microscopy

To immunostain tracheal epithelial cells, the trachea samples were isolated from mice euthanized with Avertin. The epithelium was scraped off the trachea using a scalpel in ice-cold PBS. The cell suspension was then transferred onto the coverslips coated with poly-L-lysine (P1399, Sigma) for 15 min. The epithelial cells were then fixed with freshly prepared 4% PFA in PBS for 15 min at room temperature (RT). Immunostaining of mEPCs was carried out as described (*H. Zhao et al., 2019*). In brief, mEPCs on coverslips were pre-extracted with 0.5% Triton X-100 in PBS for 40 sec, followed by fixation with 4% PFA in PBS for 15 min at RT. The epithelial cells or mEPCs on coverslips were permeabilized with 0.5% Triton X-100 in PBS for 15 min and blocked with 4% BSA in TBST (50 mM Tris, 150 mM NaCl, 0.05% Tween-20, pH 7.4) for 1 h. Primary antibodies and secondary antibodies diluted in the blocking buffer were applied to samples for 16 h at 4°C and for 1 h at RT, respectively. Confocal images were acquired with a Leica TCS SP8 system with a 60× /1.40 oil-immersion objective, and Z-stack images were generated using maximum intensity projections. 3D-SIM images were captured with a Delta Vision OMX SR imaging system (GE Healthcare) equipped with a Plan Apo 60× /1.42 oil-immersion objective lens (Olympus). Serial Z-stack imaging was performed at 125-nm intervals, and images were processed using the SoftWoRx software.

### Immunoprecipitation, GST pull-down, and immunoblotting

Cells were gently washed with 5 ml of PBS and lysed in 1 ml of pre-chilled high-salt lysis buffer (1% NP-40, 500 mM NaCl, 50 mM Hepes, 5 mM EDTA, 1 mM Na_3_VO_4_) supplemented with 1 mM PMSF and protease inhibitor (539134, Calbiochem). After centrifuging for 10 min at 4°C, the supernatant was collected and incubated with anti-GFP-agarose beads (GNA-25-500, Lalbead) at 4°C for 4 h. The beads were then thoroughly washed with high-salt lysis buffer, and the bound proteins were eluted with the SDS sampling buffer. For GST pull-down, truncated proteins were expressed using the BL21 CodonPlus (DE3) RIPL bacteria strain (230280, Agilent). GST- and His-tagged proteins were purified using glutathione-agarose beads (G4510, Sigma) and nickel-nitrilotriacetic acid (Ni-NTA) agarose beads (Qiagen, 30210), respectively. Purified GST- and His-tagged proteins were mixed in high-salt lysis buffer and incubated with glutathione-agarose beads at 4°C for 4 h. Enriched proteins were eluted in the SDS sampling buffer.

Eluted protein samples were further separated by electrophoresis and transferred to a nitrocellulose membrane (66485, Cytiva). After blocking with 5% skim milk for 1 h at RT, the membranes were placed in the TBST containing 5% bovine serum albumin (BSA; B24726, Abcone) and primary antibody overnight at 4°C. Membranes were then incubated with horseradish peroxidase (HRP) conjugated secondary antibody for 1 h at RT, and proteins were detected using the Omni-ECL efficient light chemiluminescence kit (SQ203L, Epizyme) and imaged with a Tanon 5200 imaging system.

### Electron microscopy

mEPCs serum-starved for 10 days were fixed in 2.5% glutaraldehyde (GA) and 4% PFA for 1 h at 4°C. The trachea and lateral wall of the brain were dissected from mice that had been deeply anesthetized with avertin and then transcardially perfused with PBS and 4% PFA in PBS. Tissue samples were then placed into PBS containing 2.5% GA and 4% PFA for 2 h at RT. mEPC and tissue samples were further fixed at 4°C overnight. Samples were then rinsed three times with 0.1 M phosphate buffer for 10 minutes each time and post-fixed in 1% OsO_4_ for 1 h at RT. Subsequently, they were dehydrated in a gradient of ethanol (50%, 70%, 80%, 90%) for 10 min each time and further dehydrated using absolute ethanol. For SEM, samples were dried using the critical point drying method, gold-coated by sputtering, and observed under a scanning electron microscope (TM3030, Hitachi Asia Limited) with an accelerating voltage of 15 kilovolts. For TEM, samples were embedded in Epon 812 resin, polymerized, and sectioned at a thickness of 60 nm. Ultra-thin sections were stained with 2% uranyl acetate for 10 min and 1% lead citrate for 5 min, and then imaged using an HT-7800 transmission electron microscope (Hitachi Asia Ltd.).

### Model building

The density maps of the C2 microtubule (EMD-24191) and the C1 microtubule (EMD-24207) were aligned and stitched together to reconstruct a complete C1-C2 repeating unit of the central apparatus using UCSF ChimeraX (*Han et al., 2022; Meng et al., 2023*). The pseudo-atomic models of the C2a projection were constructed by docking the *Chlamydomonas* C2a projection (PDB: 7SOM) into the corresponding region of the integrated C2 density map (*Gui et al., 2022*). The docking was performed as a rigid-body fit using the ‘Fit in Map’ tool in UCSF ChimeraX, which optimizes the correlation between the molecular model and the cryo-EM density. Using the established model, we selectively removed the C2a-specific density and the corresponding superimposed atomic model to schematically illustrate the structural consequences of the mutations in this study.

## Statistical analysis

Each experiment was conducted with a minimum of three biological repeats. For mouse experiments, at least 3 mice per genotype were used for analyses. For immunofluorescence staining, histological staining, and electron microscopy analysis, one representative image from at least 3 mice per genotype was presented. Quantitative results were presented as mean ± standard deviation (SD) unless otherwise specified in the figure legend. Statistical analyses were performed using GraphPad Prism software, and comparisons were performed using Dunnett’s one-way ANOVA to compare multiple treatment groups. Statistical significance was defined as P < 0.05.

## Data availability

All data needed to evaluate the conclusions in the paper are present in the paper and its Supplementary Materials. All materials generated in this study are available upon request.

## Acknowledgements

We thank Heng Guo (Electron Microscopy Core) and Ying Li (Light Microscopy Core) at Shandong Normal University for their assistance in imaging. This work was supported by the National Natural Science Foundation of China (32270807, 32300694, and 32470811).

## Author contributions

**Qian Lyu**: Investigation, Validation, Formal analysis; **Qingchao Li**: Investigation, Visualization; **Jingrui Li**: Validation, Funding acquisition; **Jiajun Luo**: Investigation; **Shanshan Nai**: Investigation; **Chunyu Liu**, **Hongbin Liu**, **Xueliang Zhu**: Resources; **Ting Song**: Funding acquisition, Formal analysis, Writing - Review & Editing; **Min Liu**: Conceptualization, Supervision, Writing - Review & Editing; **Huijie Zhao**: Conceptualization, Funding acquisition, Formal analysis, Supervision, Writing - Review & Editing.

## Figure supplements

**Figure 1—figure supplement 1. Mice lacking C2a proteins display phenotypes associated with PCD.** (**A**–**C**) Real-time PCR analyses showing the expression levels of *Ccdc108*, *Mycbpap*, and *Cfap70* in various mouse tissues. The expression was normalized using the corresponding *Gapdh* as the reference gene and baseline 1 (heart) as the reference sample (△△C_T_ method). Data are from three independent biological repeats and are presented as mean ± SEM. (**D**) Typical images of tracheal MCCs from WT and *Ccdc108* KO mice immunostained with the indicated antibodies. Note that the staining of CCDC108 in *Ccdc108* KO cells is invisible. (**E**) Representative images of WT and *Ccdc108* KO mice at eight weeks of age. (**F**) Representative images of serial vibratome sections of the brains from WT and *Ccdc108* KO mice at twelve weeks of age.

**Figure 2—figure supplement 1. Loss of C2a proteins leads to ultrastructural defects in the cilia axoneme.** (**A**, **B**) SEM images of ependyma (**A**) and trachea (**B**) epithelia isolated from WT, *Ccdc108* KO, *Mycbpap* KO, and *Cfap70* KO mice at two weeks of age. (**C**, **D**) TEM images and quantifications of ciliary axonemes in *Ccdc108* KO mEPCs serum-starved for 5 days. At least 50 axonemes from 3 mice (per genotype) were scored. Data are presented as mean ± SD.

**Figure 2—figure supplement 2. Ciliary motilities in representative WT, *Ccdc108* KO, *Mycbpap* KO, or *Cfap70* KO mEPCs.** Motilities of multicilia in WT, *Ccdc108* KO, *Mycbpap* KO, or *Cfap70* KO mEPCs were stained with SiR-tubulin and live imaged. Image sequences are played back at 5 frames per second.

**Figure 3—figure supplement 1. C2a proteins display mutual interactions.** (**A**–**G**) Co-IP and immunoblotting analyses in HEK293T cells exogenously expressing indicated proteins. GFP-tagged proteins in (**A**, **D**, and **E**) and HA-tagged proteins in (**B**, **C**, **F**, and **G**) were immunoprecipitated with anti-GFP and anti-HA agarose beads, respectively. Blots were probed with the indicated antibodies. Luci, luciferase.

**Figure 5—figure supplement 1. Loss of C2a proteins has no obvious impact on non-C2a ciliary proteins.** (**A**–**C**) Representative confocal images of mEPC cultured from WT, *Ccdc108* KO, *Mycbpap* KO, or *Cfap70* KO mice immunostained with the indicated antibodies. Note that the ciliary staining of SPEF1, HYDIN, and RSPH3 was unchanged in KO samples.

**Figure 6—figure supplement 1. Identification of ARMC3 and MYCBP as new C2a components.** (**A**, **B**) Co-IP and immunoblotting showing interactions between MYCBPAP and CFAP70, ARMC3, and MYCBP. Co-IP was performed with a normal guinea pig IgG and a guinea pig polyclonal anti-MYCBPAP antibody in lysates from mouse trachea and mEPCs. (**C**) Immunoblotting of mEPCs harvested at the indicated serum-starvation days for the indicated proteins. GAPDH is used as a loading control.

