## Supplementary figures and images for "Proteomic composition and mutual assembly of the C2a projection in vertebrate motile cilia"

### Figure 1-figure supplement 1

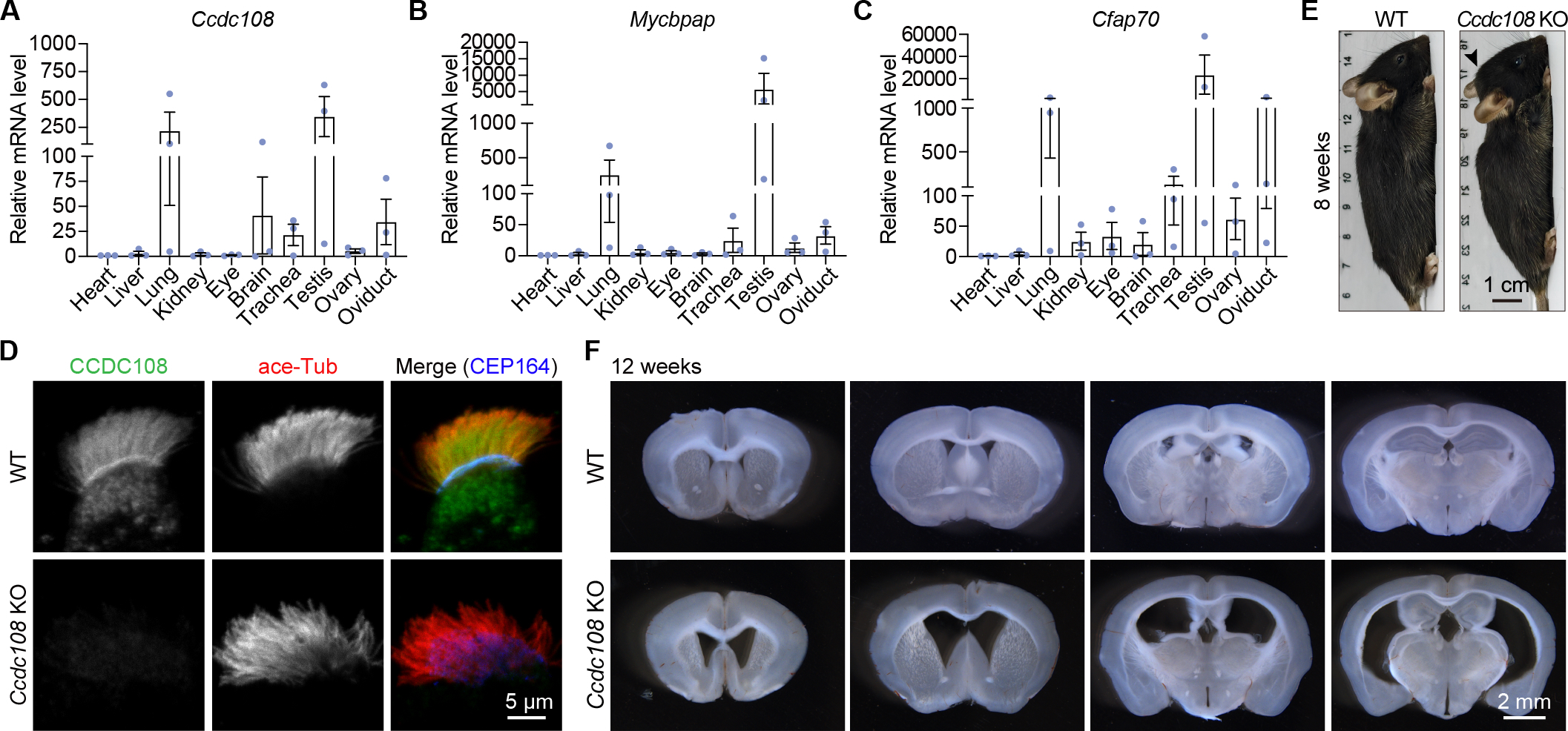

### Figure 2-figure supplement 1

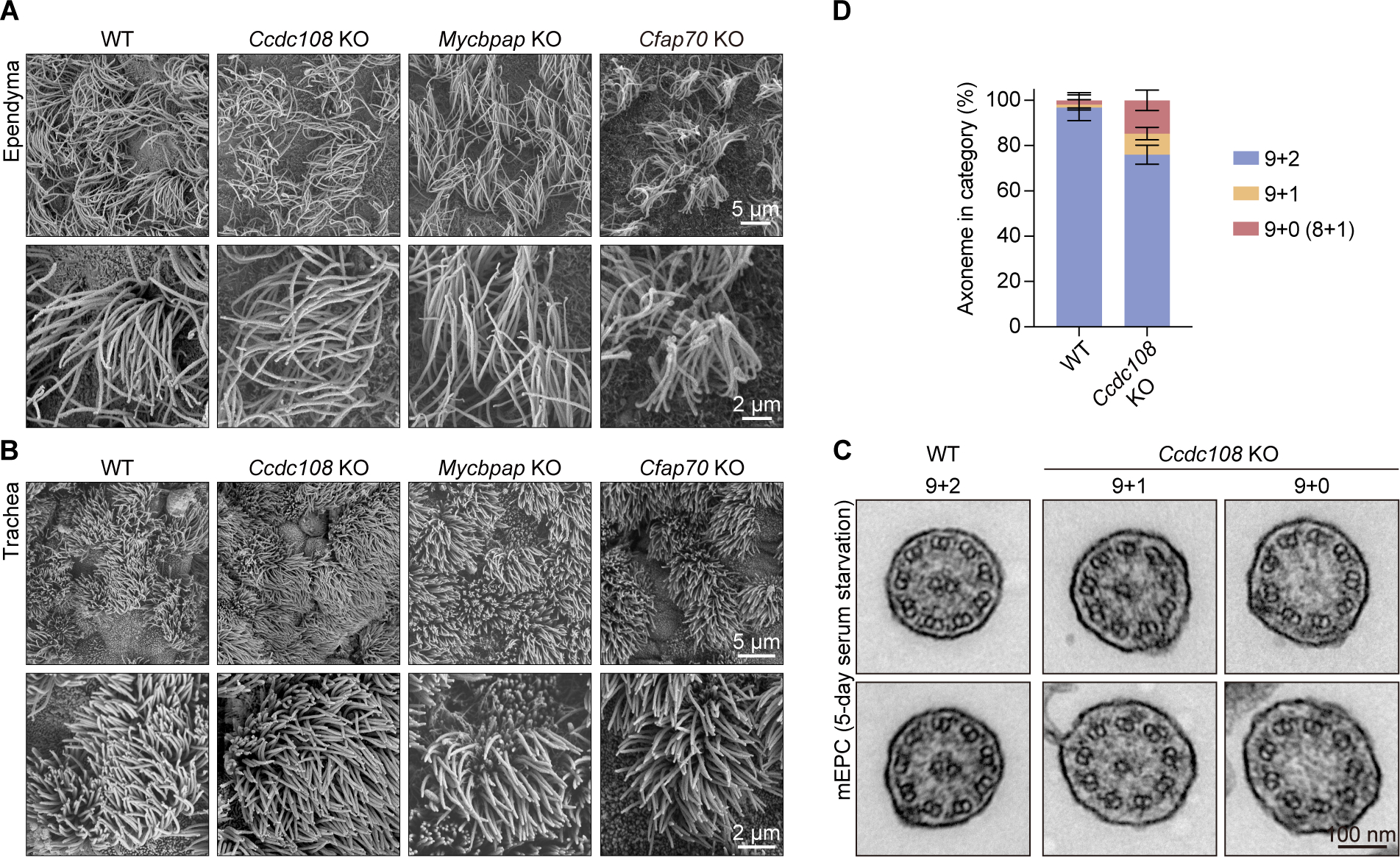

### Figure 3-figure supplement 1

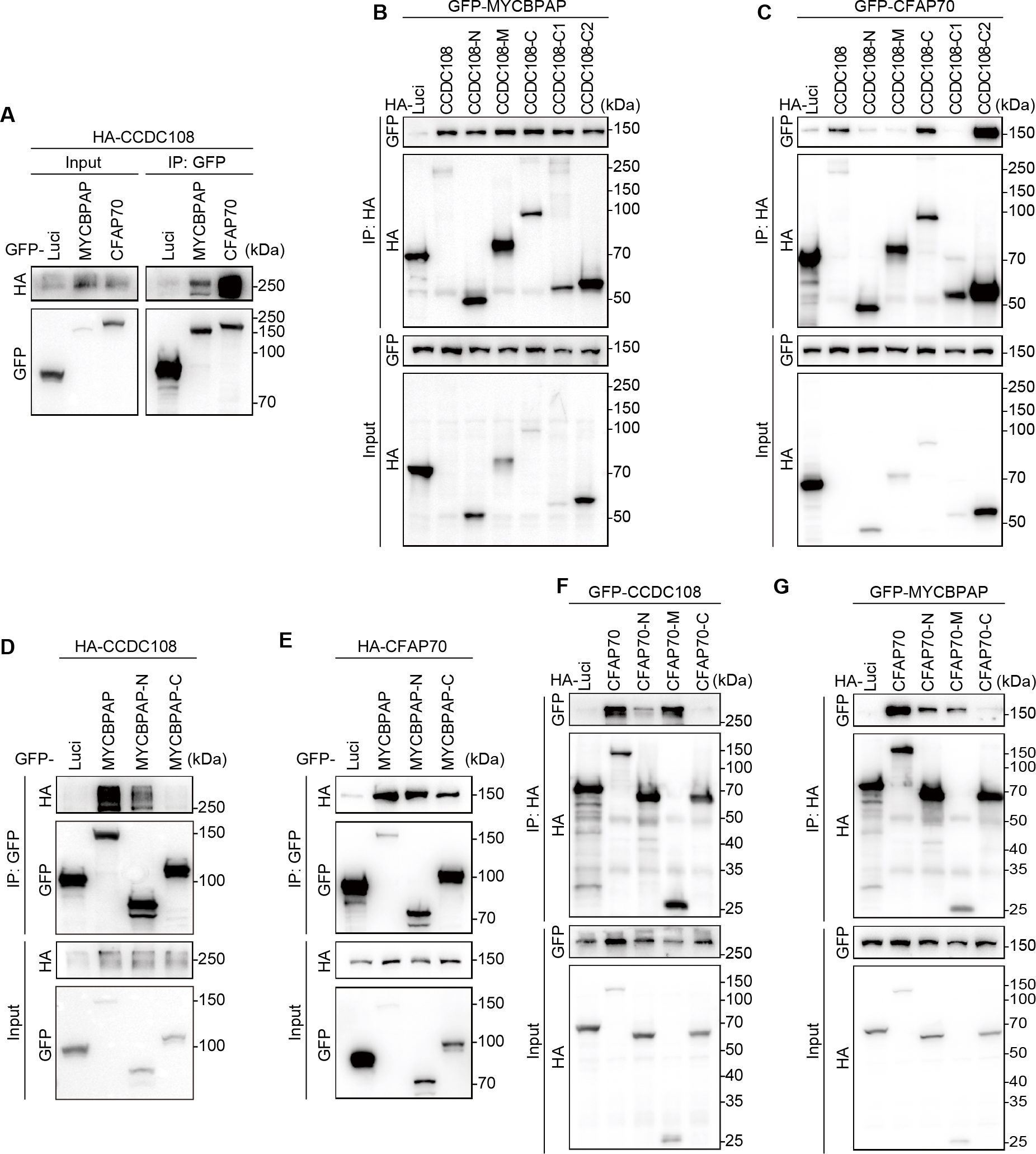

### Figure 5-figure supplement 1

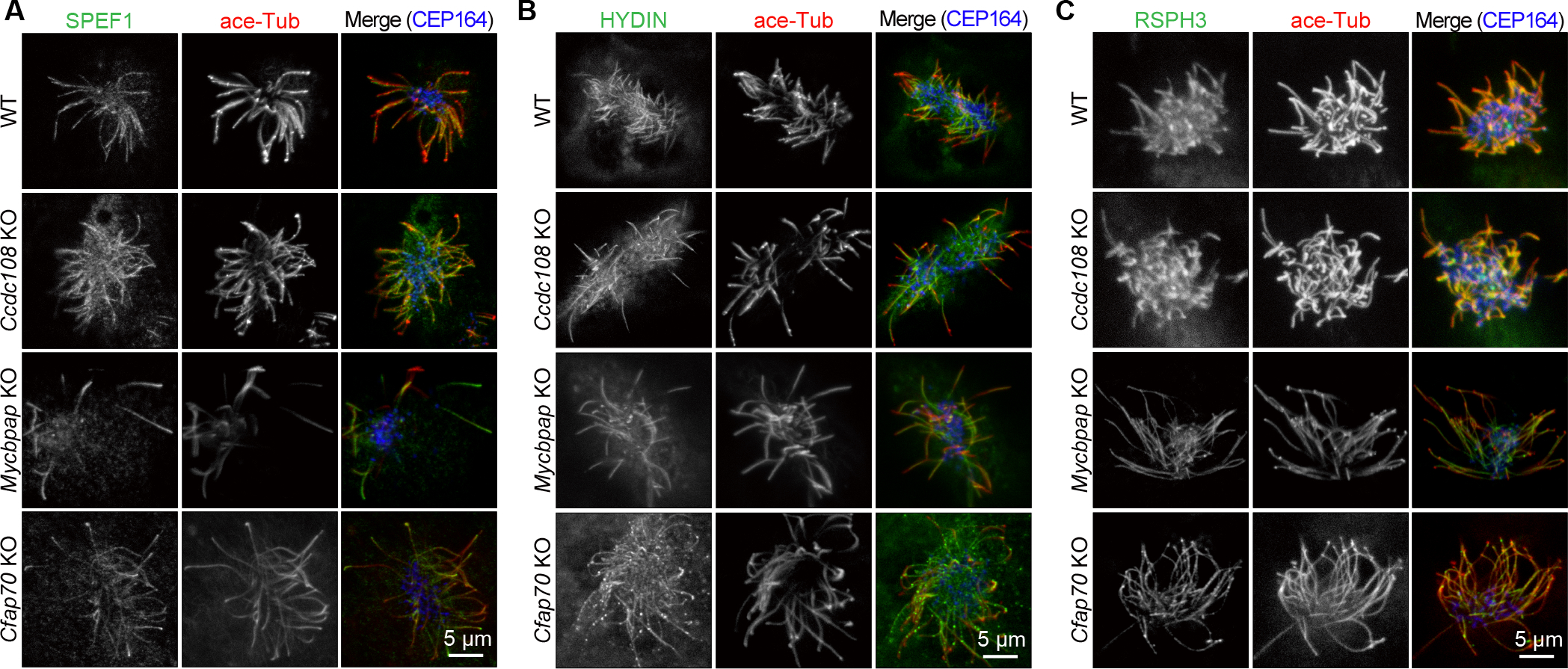

### Figure 6-figure supplement 1

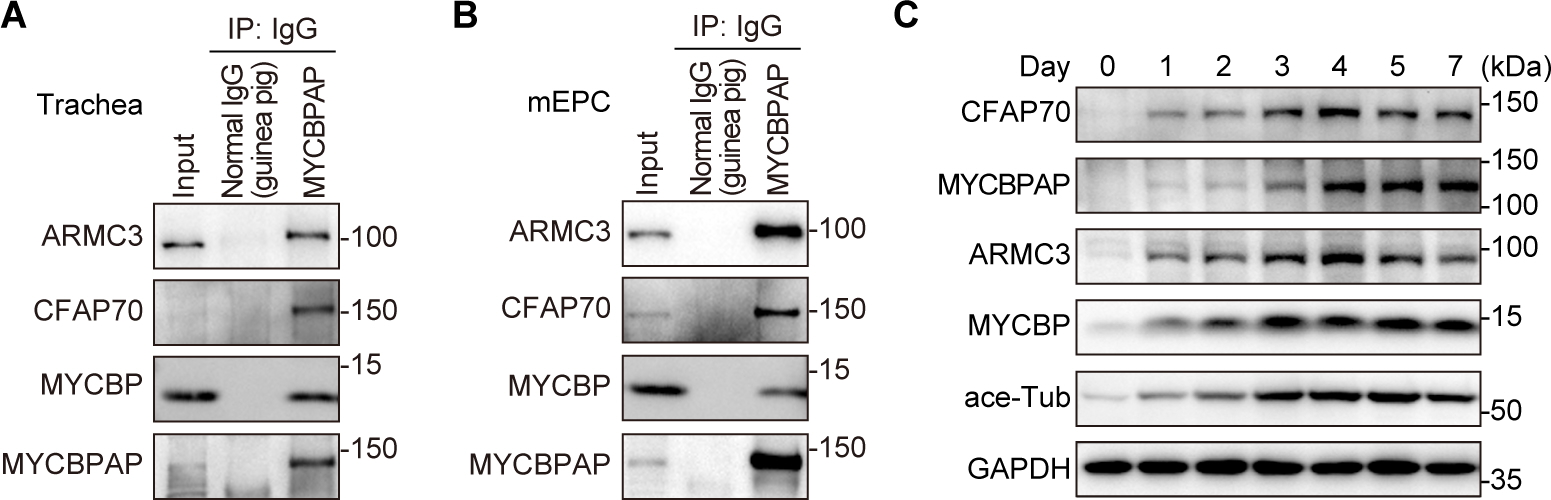
